# The origin of septin ring size control in budding yeast

**DOI:** 10.1101/2024.07.30.605628

**Authors:** Igor Kukhtevich, Sebastian Persson, Francesco Padovani, Robert Schneider, Marija Cvijovic, Kurt M. Schmoller

## Abstract

The size of organelles and cellular structures needs to be tightly controlled and adjusted to the overall cell size to ensure cell function. A prominent example of a self-assembly process forming a structure whose size scales with cell size is Cdc42-driven cell polarization and subsequent septin ring formation in *Saccharomyces cerevisiae*. Despite extensive research, the mechanisms that determine Cdc42 cluster and septin ring size are still unclear. Combining computational modeling, live-cell imaging and genetic manipulations, we show here that positive feedback in the polarization pathway, together with the amount of polarity proteins increasing with cell size, can account for the increase of the Cdc42 cluster area with cell size, and as a consequence also the scaling of the septin ring diameter. We demonstrate that in *bni1Δ* cells, which have a larger septin ring compared to wild-type cells but a similarly sized Cdc42-GTP cluster, disruption of F-actin cable assembly and polarization toward the bud site results in diffuse exocytosis, causing septin ring enlargement. Furthermore, we find that in cells with impaired negative feedback in the polarization pathway, the Cdc42-GTP cluster area expands compared to wild-type cells, while the septin ring diameter remains mostly unchanged. This decoupling can be partially explained by a high recruitment rate of septin via polarity factors. Taken together, we provide insights into the origin of septin ring size control, in particular the scaling with cell volume, by integrating new experimental findings and mechanistic models of budding yeast polarization.

## Introduction

Self-assembly processes coordinate the reproducible and timely formation of subcellular structures with a defined spatiotemporal organization and are thereby critical for cell survival. A common model for self-assembly is the process of bud site formation in the budding yeast *Saccharomyces cerevisiae*. Bud formation is a multistep process that employs regulatory strategies that are widespread across cellular processes and organisms [Bi and Park, 2012; Marquardt et al, 2019]. This includes the selection of a single bud site partially through symmetry breaking via cell polarization, followed by septin assembly and the creation of a bud neck of a specific size at the polarization site [Witte et al., 2017; Moran et al., 2019].

Cell polarization in budding yeast is well-studied experimentally and computationally [Park and Bi, 2007; Bement et al., 2024]. During polarization, Cdc42 cycles between three states (Fig. 1a): an active membrane-localized GTP-bound-state promoted by guanine nucleotide-exchange factors (GEFs), an inactive membrane-localized GDP-bound-state promoted by GTPase-activating proteins (GAPs), and a cytosolic GDI-bound state. Cdc42-driven recruitment of the GEF Cdc24 [Bose et al., 2001; Butty et al., 2002], combined with the fact that 2D diffusion of proteins in the membrane is slower than 3D diffusion in the cytosol [Woods and Lew, 2019], creates a positive feedback system that promotes the formation of a single active Cdc42-GTP cluster. In support of the central role of positive feedback, optogenetic recruitment of the GEF Cdc24 promotes polarization at the recruitment site [Witte et al, 2017]. Furthermore, computational models of positive feedback capture several observed phenomena such as competition between polarity clusters [Goryachev and Leda, 2017].

**Figure 1.**
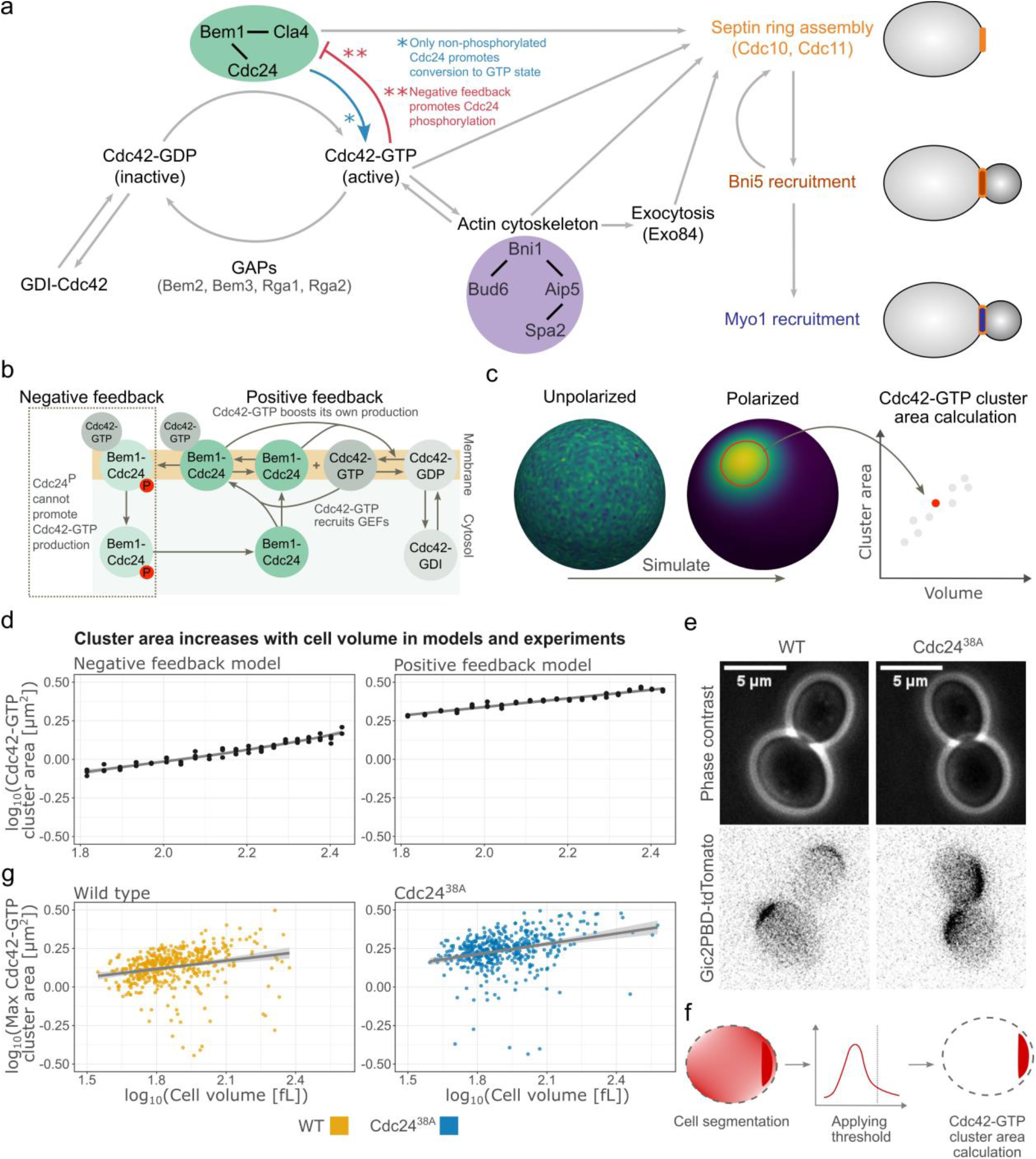
Regulatory pathways and computational modeling of Cdc42 polarization. (a) Schematics of the regulatory pathways controlling cell polarization and downstream contractile ring assembly at the polarization site. (b) Schematic of the positive and negative feedback models for Cdc42-GTP polarization. In the positive feedback model, the components inside the dashed box are removed. P denotes phosphorylation. (c) Starting from a randomly perturbed unstable steady state, the model was simulated to a polarized steady state, and the Cdc42-GTP cluster area was then determined. (d) Cdc42-GTP cluster area measured at steady-state (after long simulation time) for the model plotted against cell volume in double logarithmic scale. In each case, for n=60 (three replicates per volume) cells with volumes in the range of 65 to 270 fL were simulated starting from random initial conditions. Left plot: Negative feedback model, slope = 0.35 (p < 2*10^−16^). Right plot: Positive feedback model, slope = 0.33 (p < 2*10^−16^). (e) Representative microscopy images of budding yeast cells (phase contrast) and Cdc42-GTP clusters (Gic2PBD-tdTomato) for WT and Cdc24^38A^ (negative feedback mutant) cells. (f) Cdc42-GTP cluster area is measured from microscopy images by applying a threshold and measuring the maximum cluster area reached before bud emergence (for details, see Methods). (g) Maximum Cdc42-GTP cluster area measurements from microscopy images plotted against corresponding cell volume in double logarithmic scale. Left: WT (n = 463 cells), slope = 0.18 (p < 2.6*10^−7^). Right: Cdc24^38A^ (n = 478), slope = 0.39 (p < 2*10^−16^).

In addition to positive feedback, negative feedback within the polarization pathway has been proposed [Kuo et al., 2014; Rapali et al., 2017]. In a phosphosite Cdc24 mutant, excess Cdc42 accumulates at the bud site [Kuo et al., 2014], suggesting a major feedback operates via inhibitory phosphorylation of the GEF Cdc24. Furthermore, *in vitro* experiments have shown that Bem1’s ability to activate Cdc24 is reduced when Cdc24 is phosphorylated [Rapali et al., 2017]. Even though the precise mechanisms of negative feedback remain unknown, computational modeling has shown that it enhances the robustness of the polarization [Howell et al., 2012], suggesting it is one of the multiple mechanisms ensuring reliable polarization.

When the Cdc42-GTP cluster reaches its maximum size, septin recruitment starts, followed by septin ring assembly [Park and Bi, 2007]. Previously, we demonstrated that the diameter of the budding yeast septin ring increases with cell volume [Kukhtevich et al., 2020]. We also found that the Cdc42-GTP cluster area increases with cell volume and that the septin ring diameter can be decoupled from the cluster area by deleting the formin *BNI1* [Kukhtevich et al., 2020]. Specifically, the septin ring diameter is larger in *bni1Δ* compared to wild-type (WT) cells, even though the Cdc42 cluster area is largely unchanged. While this suggests that the size of the septin ring is set through a combination of septin recruitment and Cdc42 polarization, the mechanisms underlying size homeostasis of the structures at the budding yeast bud neck are still elusive.

Here, we use computational modeling in combination with yeast genetics and microfluidics-based time-lapse imaging to investigate the origin of the cell-volume dependence of the Cdc42 cluster area and septin ring diameter. We find that positive feedback in the polarization pathway, together with the amount of key polarity proteins increasing with cell volume, can account for the increase of Cdc42-GTP cluster area with cell size. As a consequence, the septin ring diameter then also increases with cell size. In *bni1*Δ cells, F-actin cable assembly and polarization toward the bud site are disrupted, resulting in diffuse exocytosis. Moreover, our simulation results show that if exocytosis supports septin recruitment, diffused exocytosis can explain the increased septin ring diameter despite unchanged Cdc42-GTP cluster area. Consistent with experimental findings, our model also predicts that disruption of the Cdc24-dependent negative feedback in the polarization pathway leads to an increased Cdc42-GTP cluster area. Despite this increase in the Cdc42-GTP cluster area, the septin ring diameter is largely unaffected. This decoupling between septin ring diameter and Cdc42 cluster area can be partially recapitulated in our model by increasing the rate at which septin is recruited by polarity factors.

## Results

### Negative feedback in the polarization pathway is not required for scaling of the Cdc42-GTP cluster area with cell volume

Given the key role of Cdc42-GTP in bud site selection and initiation of septin ring assembly, we first investigated Cdc42 polarization and subsequently the consequences on septin ring formation.

We have previously shown that the Cdc42-GTP cluster area increases with cell size [Kukhtevich et al. 2020]. To gain insights into the mechanisms underlying this scaling, we used computational modeling. As Cdc42 polarization is driven by a positive feedback loop [Kozubowski et al., 2008; Witte et al., 2017], and also likely a negative feedback loop [Kuo et al., 2014], we implemented a previously published model with both a positive and negative feedback [Kuo et al., 2014]. In this model, Cdc42-GTP attracts its own effectors (Bem1-Cdc24-complex) to form a Bem1-Cdc24-Cdc42 complex, which in turn activates Cdc42 (Fig. 1b). Combined with membrane diffusion being slower than cytosolic diffusion, this creates a positive feedback system where Cdc42 can polarize from stochastic fluctuations [Goryachev and Pokhilko, 2008; Wu et al., 2015]. Moreover, to account for reported phosphorylation-driven negative feedback [Kuo et al., 2014], it is assumed in the model that when the Bem1-Cdc24-Cdc42 complex reaches a high concentration, the Bem1-Cdc24 complex component is phosphorylated. This inhibits the ability of Bem1-Cdc24 to activate Cdc42 (Fig. 1b left). We refer to this model as the ‘negative feedback model’. For simulations, we approximate the yeast cells as spheres (Fig. 1c), and we simulate the model over a high-density finite-element (FEM) mesh.

Starting from a steady state with random perturbations to facilitate polarization, we simulated 60 cells undergoing the transition from an unpolarized to a polarized state with cell volumes ranging from 65 to 270 fL. We then determined the Cdc42-GTP cluster area for each cell at the final steady-state time point (Fig. 1c). In line with experimental observations [Kukhtevich et al., 2020], we find that in the negative feedback model, the cluster area increases with cell volume (R^2^=0.95) (Fig. 1d left), consistent with a power law relationship *A ∼ V*^*0.35*^.

Next, we aimed to understand the contributions of the negative and positive feedbacks in the model and asked whether the positive feedback is sufficient for the Cdc42-GTP cluster area scaling. To assess this, we removed the negative feedback in the model (Fig. 1b left). We refer to this model as the ‘positive feedback model’. Again, cluster area followed a power law relationship with cell volume, *A ∼ V*^*0.33*^ (R^2^=0.98) (Fig. 1d right). The cluster area is also larger compared to the negative feedback model (Fig. 1d). To validate our results beyond this single model, we applied a similar simulation approach to an alternate positive feedback polarization model [Borgqvist et al., 2021] and again found that the Cdc42-GTP cluster area increases with cell volume (Fig. S1). Additionally, since the mechanisms governing the negative feedback are not fully elucidated, we also explored an alternative way to simulate the effect of decreasing Cdc42-GTP activation. Specifically, we increased GAP activity in the positive feedback model. Consistent with the negative feedback model, also with increased GAP activity cluster area scaled with cell volume, and the cluster area was smaller (Fig. S2a-b)

Our simulation results suggest that the positive feedback alone is sufficient to establish the increase of the Cdc42-GTP cluster area with cell volume. To validate these simulations experimentally, we investigated a strain in which the negative feedback loop is disrupted by a point mutation in Cdc24 [Kuo et al., 2014]. Briefly, Cdc42-GTP activates the p21-activated kinase Cla4, which then phosphorylates Cdc24. In the case of a Cdc24^38A^ phosphosite mutant strain [Kuo et al., 2014], phosphorylation is prevented, weakening the negative feedback while keeping the positive feedback intact. To measure the Cdc42-GTP cluster area, we used a microfluidic device previously described in [Kukhtevich et al., 2022] in combination with time-lapse imaging (Fig. 1e, f). As predicted by the model, the maximal Cdc42-GTP cluster area prior to bud emergence increased with cell volume for both Cdc24^38A^ and WT strains (Fig. 1g). To confirm this result, we also employed a previously established system based on the tunable expression of the cell size regulator Whi5 to increase the range of experimentally accessible cell volumes [Kukhtevich et al., 2020; Claude et al., 2021]. With this approach, we again found that cluster area increases with cell volume for both wild-type Cdc24 and Cdc24^38A^ (Fig. S3). Taken together, this confirms the model prediction that the positive feedback alone qualitatively accounts for the scaling of Cdc42-GTP cluster area with cell volume.

### Scaling of Cdc42-GTP cluster area with cell volume requires that the amount of polarity proteins increases with cell volume

Our simulation results revealed that the positive feedback is sufficient for the Cdc42-GTP cluster area to increase with cell volume (Fig. 1). However, this is under the assumption that the concentration of all polarity proteins in the model is constant. While global protein amounts increase with cell volume, this increase is not proportional for every protein, leading to a cell-size-dependent decrease in the concentration of specific proteins [Swaffer et al., 2021; Lanz et al., 2024]. To test how the Cdc42 cluster area would be affected by a dilution of polarity proteins with cell volume, we simulated the model for different dependencies of the concentration of polarity proteins on cell volume. To characterize this dependency, we used the slope of the double-logarithmic dependence of protein concentration on cell volume [Claude et al., 2021; Lanz et al., 2024]. Specifically, a ‘protein slope’ equal to –1 is equivalent to concentrations that decrease in inverse proportion with cell volume, and a slope equal to 0 means that concentration is constant with volume (Fig. 2a). We simulated n=60 cells (65 to 270 fL) under three conditions: i) constant protein amounts (protein slope = –1, Fig. 2a, b), and ii-iii) two different scenarios of amounts that increase sublinearly with cell volume, leading to a weaker cell-size-dependent decrease of the corresponding concentrations (protein slope = –0.2 or –0.44, respectively, Fig. 2b). The GAP concentration was kept constant in all simulations. If in the negative feedback model (WT model), the protein amount increases, cluster area also increases with cell volume (protein slope = –0.2 or –0.44, Fig. 2c). Similar trends hold for the positive feedback (Cdc24^38A^) model (Fig. 2c). However, when Cdc42 and Bem1-Cdc24 complex amounts were constant, the negative feedback model did not polarize when we increased volume. In the positive feedback model, the Cdc42-GTP cluster area did not scale with cell volume (protein slope = –1, Fig. 2c).

**Figure 2.**
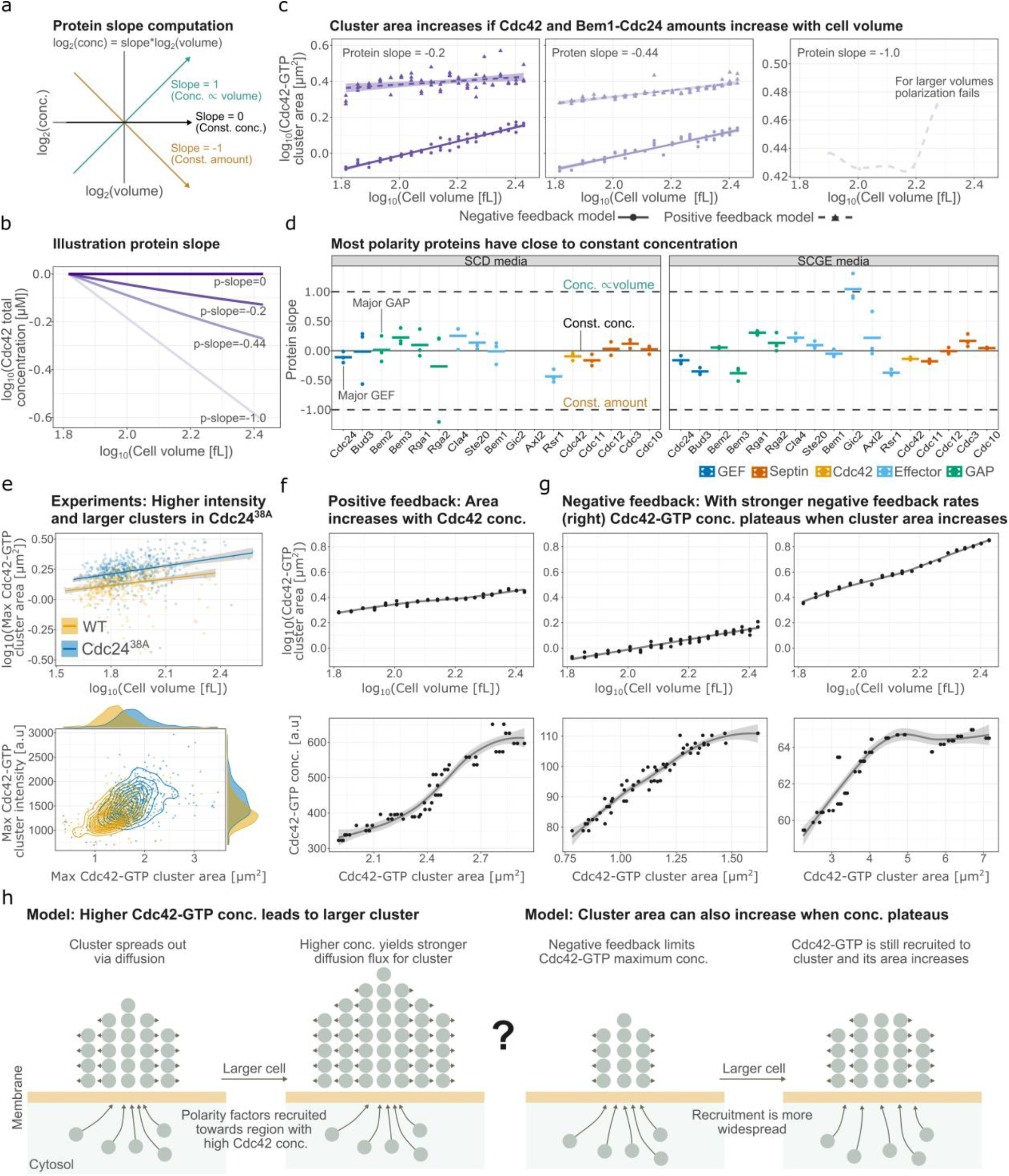
The amount of key polarity proteins increases with cell volume to ensure robust polarization, while the interplay between positive and negative feedback determines the Cdc42-GTP concentration and cluster area. (a) Illustration of how concentration changes with cell volume for different protein slopes. (b) Simulated dependence of Cdc42 concentration on cell volume for the different protein slopes. (c) Modeling results for Cdc42-GTP cluster area measured at steady-state (after long simulation time) plotted against cell volume in double logarithmic scale for the following conditions: Cdc42 and Bem1 complex amount increase with cell volume (protein slope = –0.2 or –0.44), Cdc42 and Bem1 complex amounts constant (protein slope = –1). The two left panels show results from the negative and positive feedback model; only the positive feedback model is shown in the right panel as it was the only model to polarize for this condition. In each case, n=60 cells (three replicates per volume) with volumes ranging from 65 to 270 fL were simulated starting from random initial conditions. For the two leftmost panels, solid lines show linear regression fits. (d) Protein slopes for the majority of polarity proteins are close to 0, implying constant concentrations in SCD or SCGE media. Data from [Lanz et al., 2024]. Bars show the mean value of n=3 biological replicates. (e) Experimental results for WT and Cdc24^38A^ cells: (top) maximum Cdc42-GTP cluster area prior to bud emergence plotted against corresponding cell volume in double logarithmic scale, (bottom) Maximum Cdc42-GTP intensity in the cluster at the time where the cluster reaches its maximum area, plotted against the corresponding Cdc42-GTP cluster area. WT: n = 463 cells; Cdc24^38A^: n = 478 cells. Solid lines in the top plot show linear regression fits. (f) Results obtained with the positive feedback model: (top) Cdc42-GTP cluster area measured at steady-state (after long simulation time) plotted against cell volume in double logarithmic scale, (bottom) Cdc42-GTP concentration at the same time-point as cluster area was measured, plotted against the Cdc42-GTP cluster area. n=60 (three replicates per volume) cells with volumes ranging from 65 to 270 fL were simulated starting from random initial conditions. Lines show loess smoothings. (g) Results for the negative feedback model are shown as in panel b. The left plot corresponds to normal phosphorylation rates in the model; in the right plot, both the phosphorylation and dephosphorylation rates of the Bem1-Cdc24 are increased. As in panel f, n=60 cells, and the lines show loess smoothings. (h) Illustration of how higher Cdc42-GTP concentration in the cluster can lead to an increase in its area (left), and how the cluster area can increase even when the Cdc42-GTP concentration plateaus in the presence of negative feedback (right).

Our modelling thus predicts that for the Cdc42-GTP cluster area to increase with cell volume, the amount of polarity proteins must increase with volume. To investigate if this is indeed the case, we analyzed a recent proteomics dataset where a triple-SILAC workflow was used to measure protein concentrations of different-sized cells that were obtained through G1 arrest in either glucose (SCD) or ethanol/glycerol-containing media (SCGE) [Lanz et al., 2024]. To characterize the cell size dependence, a protein slope was then computed for each polarity protein (Fig. 2d). We found that most polarity proteins are maintained at close to constant concentrations across cell volumes (protein slope = 0 in Fig. 2d), which is in line with an earlier report [Chiou et al, 2021]. Notably, the key polarity proteins Cdc42 (SCD: mean slope = –0.14, SCGE: mean slope = –0.09), Cdc24 (SCD: slope = –0.16, SCGE: slope = –0.11), Bem2 (SCD: slope = 0.05, SCGE: slope = 0.01), and Bem1 (SCD: slope = –0.05, SCGE: slope = –0.01) exhibit protein slopes close to zero, well within the range in which the model predicts the Cdc42-GTP cluster area to increase with cell volume (Fig. 2c). Similar trends hold for the polarity proteins in an experiment where different sized cells were obtained via mutations (Fig. S2c).

In summary, our computational results suggest that while the scaling of the Cdc42-GTP cluster area with cell volume is robust to weak protein dilution, the amount of key polarity proteins must increase with cell volume for the Cdc42-GTP cluster area to increase. Indeed, reanalysis of experimental data from Lanz et al. [Lanz et al., 2024] confirmed that the amount of key polarity proteins increases with cell volume.

### Potential mechanisms underlying the scaling of Cdc42-GTP cluster area with cell volume

Our experiments and modelling show that cluster area increases with cell volume (Fig. 2e). Next, we sought to identify the origin of the cluster scaling using modeling.

Diffusion is driven by concentration gradients. In both the positive and negative feedback models, the maximum concentration of Cdc42-GTP increases with cell volume (Fig. 2f, 2g left). This suggests that the Cdc42-GTP cluster area increases with cell volume because the diffusion-driven flux gets stronger due to higher concentration in the cluster center (Fig. 2h left). Supporting this idea, we found that reducing the GAP activity in the positive feedback model concentrates more Cdc42-GTPs in the center of the cluster, and also results in an increased cluster area (Fig. S2). Furthermore, in Cdc42^38A^ cells, in which the maximum intensity in the cluster based on the Gic2PBD-tdTomato reporter for Cdc42-GTP is higher than in WT cells, the cluster area also increased (Fig. 2e). Similar results hold for Whi5-induced cells (Fig. S3b, c). However, we note that in addition to the concentration of Cdc42-GTP, the absolute expression level of the reporter may affect the cluster intensities, which makes it difficult to interpret intensity differences between strains.

Interestingly, with negative feedback in the model, an increase in the Cdc42-GTP cluster area can even be observed without an increase in the Cdc42-GTP concentration. Specifically, if both the phosphorylation and dephosphorylation rates of Bem1-Cdc24, which are associated with the negative feedback, are increased sufficiently, the maximum Cdc42-GTP concentration plateaus at larger cell volumes while the cluster area still increases (Fig. 2g right). Effectively, the negative feedback restricts Cdc42 from concentrating above a certain threshold, but owing to the positive feedback, polarity proteins are recruited towards the cluster. Thus, with increased cell volume, more proteins are recruited, and cluster area increases (Fig. 2h right).

### Deletion of polarisome complex components leads to increased septin ring diameter

After successfully modelling the scaling of the Cdc42-GTP cluster area as described above, we then set out to better understand the subsequent assembly of a septin ring. In our previous work [Kukhtevich et al., 2020], we showed that similar to the Cdc42-GTP cluster area, also the septin ring diameter increases with cell volume. Interestingly, we also found that despite an unchanged Cdc42 cluster area, *bni1*Δ cells show an increased septin ring diameter compared to WT cells after accounting for cell volume (∼28% increase). This highlights that the septin ring diameter is not solely determined by the Cdc42 cluster size. Before developing a mechanistic model for septin ring assembly, we therefore decided to obtain a better understanding of how the formin and component of the polarisome complex Bni1 contributes to septin ring formation. First, we asked whether similar to *bni1*Δ, also deletion of the second yeast formin Bnr1 [Park and Bi, 2007; Yu et al., 2011] or the additional polarisome components Spa2 and Bud6 [Liu et al., 2010; Xie et al., 2019] would lead to an increased septin ring diameter. We introduced the corresponding deletions into a strain carrying an mCitrine-tagged allele of the septin Cdc10 and performed microfluidics-based time-lapse experiments. We found that *spa2Δ* and *bud6Δ* cells partially phenocopy the effect of *bni1Δ* cells on cell geometry, cell volume and Cdc10 ring diameter, which is increased by ∼13 % compared to WT cells. By contrast, the septin ring diameter is largely unaffected in *bnr1*Δ cells (Fig. 3). Based on this, we can conclude that polarisome components are more important for setting the septin ring diameter than the second formin Bnr1, and that Bni1 has a larger effect on the ring diameter than Spa2 and Bud6.

**Figure 3.**
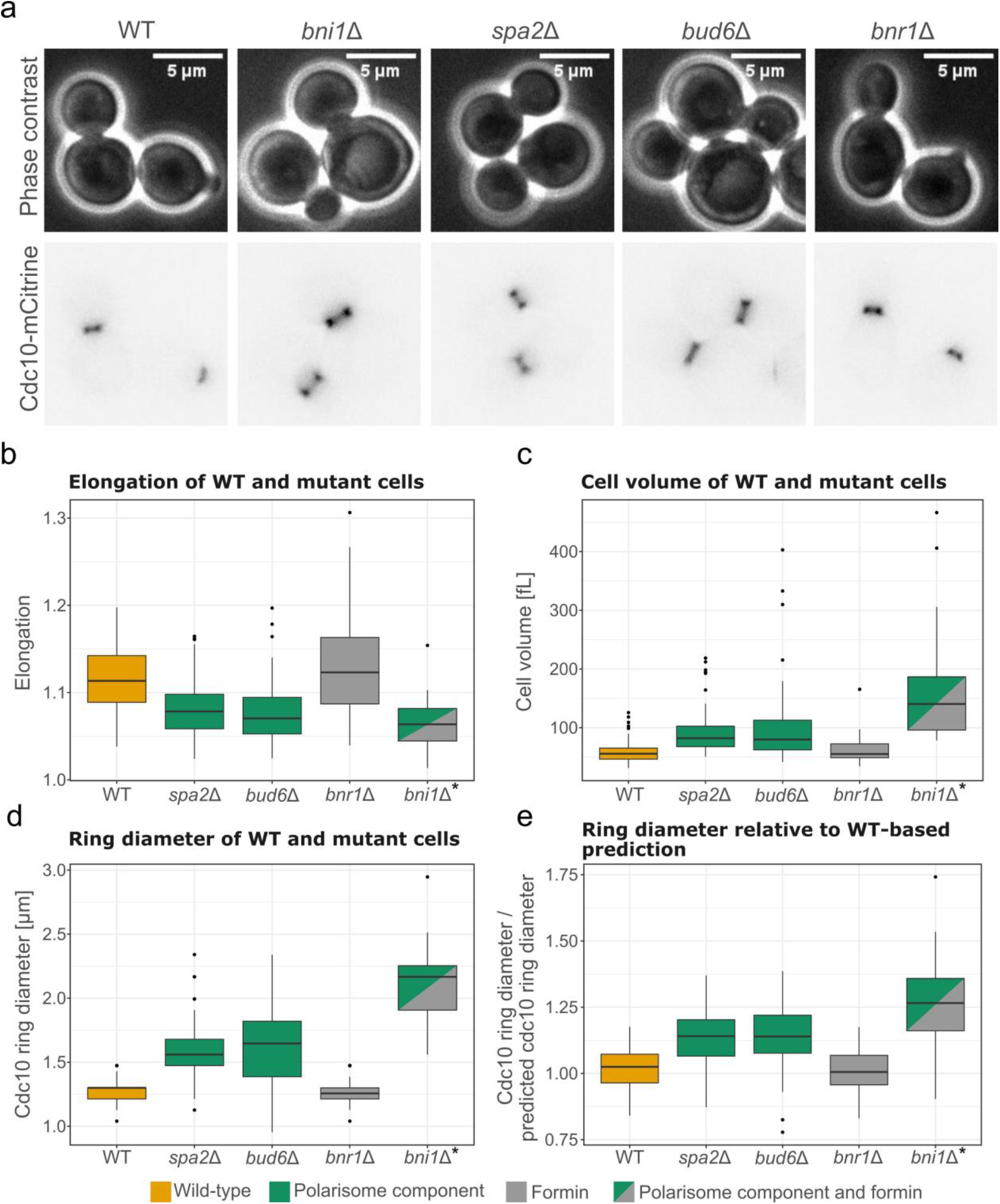
Impact of formins and components of the polarisome complex on the septin ring diameter. (a) Representative microscopy images of budding yeast cells (phase contrast) and septin ring (Cdc10-mCitrine) for WT, *bni1Δ, spa2Δ, bud6Δ* and *bnr1Δ* cells. (b) Elongation (major cell axis/minor cell axis) of WT (n = 81 cells), *spa2Δ* (n = 90 cells), *bud6Δ* (n = 83 cells), *bnr1Δ* (n = 67 cells) and *bni1Δ\** (n = 35 cells) (c) Mother cell volume for the same cells as in b. (d) Median septin ring diameter during its presence for the same cells as in panel b. (e) The predicted septin ring diameter for the same cells as in panel b was calculated by dividing the septin ring diameter of a given cell by a model prediction based on the WT data in [Kukhtevich et al., 2020]. Note that the WT dataset shown here is not the same as the data the model prediction is based on. (b-e) * data from [Kukhtevich et al., 2020].

### Spatial confinement of exocytosis is critical for septin ring diameter control

The formin Bni1 is actively involved in F-actin cable assembly [Yu et al., 2011; Liu et al., 2010; Xie et al., 2019]. Therefore, we decided to focus next on the role of F-actin in setting the septin ring diameter. To analyze F-actin cables along with the septin ring, we tagged Cdc10 with mNeptune2.5, and Abp140, which has previously been used for actin cytoskeleton visualization [Buttery et al., 2007], with mCitrine. To test a wide range of cell volumes, we again used inducible Whi5. Microfluidics time-lapse experiments revealed that in *bni1Δ* cells, F-actin cables mostly do not polarize toward the bud side, while in WT cells, cable polarization is prominent (Fig. 4a). Our analysis showed that in WT cells, the Abp140 cluster area scales with cell volume (Fig. 4b). We also observed a clear correlation between the Cdc10 ring diameter and the Abp140 cluster area (Fig. 4c). By contrast, in *bni1Δ* cells, only ∼11% of cells show clearly detectable Abp140 clusters. This suggests that F-actin cables play an important role in setting the septin ring diameter.

**Figure 4.**
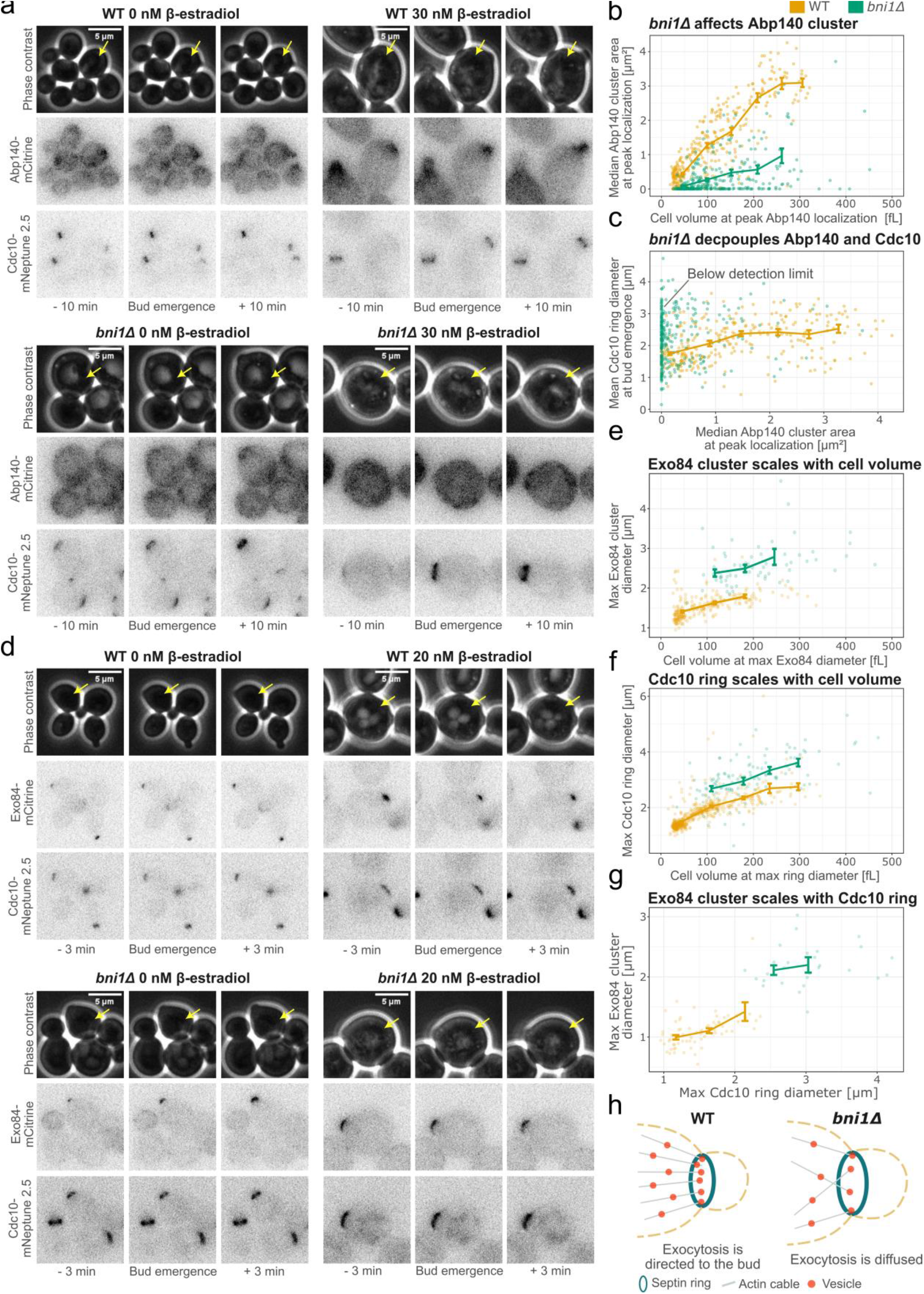
Exocytosis is diffused in *bni1Δ* cells due to disturbed F-actin cable assembly and polarization toward the bud site. (a) Representative microscopy images of budding yeast cells (phase contrast), F-actin (Abp140-mCitrine) and septin ring (Cdc10-mNeptune2.5) for WT and *bni1Δ* cells. Different cell sizes are obtained by different β-estradiol concentrations (0 or 30 nM) to induce *WHI5* expression (see methods). (b) Abp140 cluster area measurements from microscopy images plotted against corresponding mother cell volume in double logarithmic scale for WT (n = 324 cells) and *bni1Δ* (n = 319 cells). (c) Cdc10 ring diameter at bud emergence plotted against Abp140 cluster area for WT (n = 290 cells) and *bni1Δ* (n = 298 cells). (d) Representative microscopy images of cells (phase contrast), exocytosis (Exo84-mCitrine) and septin ring (Cdc10-mNeptune 2.5) for WT and *bni1Δ* cells. Different cell sizes are obtained by different β-estradiol concentrations (0 or 20 nM) to induce *WHI5* expression. (e) Exo84 cluster diameter measurements from microscopy images plotted against corresponding mother cell volume in double logarithmic scale for WT (n = 218 cells) and *bni1Δ* (n = 62 cells). (f) Maximum Cdc10 ring diameter plotted against corresponding mother cell volume in double logarithmic scale for WT (n = 556 cells) and *bni1Δ* (n = 113 cells). (g) Exo84 cluster diameter measurements from microscopy images plotted against maximum Cdc10 ring diameter for WT (n =68 cells) and *bni1Δ* (n = 28 cells). (a,d) Annotation of bud emergence corresponds to the cells highlighted with yellow arrows. (b,c,e,f,g) Solid lines show binned means and error bars show standard error. h) Suggested mechanism for septin ring enlargement in *bni1Δ* cells.

We next asked why correct F-actin cable assembly and polarization toward the bud site are important for controlling the septin ring diameter. Okada et al. [Okada et al., Dev. Cell., 2013] previously suggested that the septin ring is sculpted by polarized exocytosis, which displaces the accumulating septins at the center of the bud site and thereby relieves inhibition of Cdc42 which is mediated via septin-recruited GAPs. Since cargo vesicles are delivered along F-actin cables, actin has a direct impact on exocytosis [Park and Bi, 2007; Liu et al., 2010]. Thus, we decided to investigate if the septin ring diameter depends on exocytosis and how this dependency is affected by deleting *BNI1*. We constructed new strains in which we tagged Exo84, one of the exocyst complex subunits [Jose et al, MBoC, 2015], with mCitrine, and Cdc10 with mNeptune2.5. Again, we used inducible Whi5 to increase the range of cell volumes and performed time-lapse experiments, where we quantified the Exo84 cluster diameter together with the Cdc10 ring diameter. We found that the Exo84 cluster diameter scales linearly with cell volume on a double logarithmic scale. In addition, in *bni1Δ* cells, the Exo84 cluster diameter is larger compared to WT cells (Fig. 4d, e). Moreover, the Cdc10 ring diameter is larger in *bni1Δ* cells and scales with the Exo84 cluster diameter in both WT and *bni1Δ* cells (Fig. 4f, g). Both strains appear to follow the same scaling relationship, indicating a mechanistic link between exocytosis and septin ring diameter.

In summary, our results suggest that in *bni1Δ* cells, perturbed F-actin assembly and polarization lead to diffused, *i.e*. less focused, exocytosis, which in turn leads to an enlarged septin ring (Fig. 4h).

### Exocytosis-aided recruitment of septins explains the increase of septin ring size upon diffused exocytosis

Our results suggest that in *bni1*Δ cells, exocytosis at the bud site is more diffused than in WT cells (Fig. 4). Given that exocytosis displaces proteins from the polarity site in *S. pombe* [Gerganove et al., 2021], and that it has been suggested to sculpt the septin ring in *S. cerevisiae* by displacing septins [Okada et al., 2013], we hypothesized that diffused exocytosis causes a larger ring diameter by septin displacement (Fig. 4h). To test this, we turned to computational modeling.

Initially, we used a model inspired by Okada et al. [Okada et al., 2013], where the septin ring is sculpted by exocytosis. However, likely because this model relies on exocytosis as the sole mechanism driving ring formation, we found that it is not robust to changes in cell volume (see supplementary information). Therefore, we implemented a new model (Fig. S4a, b), which we refer to as the *Septin Binding and Exocytosis* (SBE) model.

Because septin recruitment is likely cell-cycle triggered [Lai et al., 2018], and to reduce simulation time, we decided to temporally separate Cdc42 polarization and septin ring formation in the model. Our SBE model, therefore, consists of a Cdc42 polarization module and a septin ring module (Fig. S4). In the polarization module, we used only the positive feedback model. This is because, as we demonstrated above, a positive feedback is capable of recapitulating the scaling of the Cdc42-GTP cluster area with cell volume (Fig. 1, S1), and such models have captured other behaviors such as competition between polarity sites [Goryachev and Pokhilko, 2008]. Moreover, this allows us to reduce simulation runtime for the analysis of septin ring formation (one simulation takes >100h), as the positive feedback model is faster to simulate than the negative feedback model. For the septin module, to capture that septin is recruited by and interacts with polarity factors such as Gic1/2, Axl2, Cdc24 and potentially Cdc42 itself [Iwase et al., 2006; Sadian et al., 2013; Chollet et al., 2020; Kang et al., 2024], we included the key player Axl2 as a representative for all recruitment factors. Furthermore, as Cdc24 binds Cdc11 in the Cdc42-GTP cluster but not in the septin ring [Chollet et al., 2020], and Cdc42 inhibits septin polymerization *in vitro* [Sadian et al., 2013], we model that septin binds to polarity factors, represented by Axl2, and that septin cannot polymerize when bound. This drives septin polymerization and subsequently ring formation at the cluster periphery. As a result, the septin ring can form even in the absence of exocytosis (Fig. S4e), in line with observations that ring formation can start without a clear Sec4 (exocytosis marker) signal [Lai et al., 2018]. Furthermore, exocytosis was modeled to be directed towards the Cdc42 cluster, where for the meshed model geometry, vesicles could hit a tunable percentage of the mesh nodes, and when occurring, exocytosis displaces proteins. Lastly, following earlier observations [Okada et al., 2013], septin recruits GAP proteins (Fig. S4a).

To simulate the septin ring module, we initiated the septin module of the SBE model simulations from a Cdc42-GTP cluster obtained with the Cdc42 polarization module (Fig. S4). Given this approach, the SBE model can form a septin ring, but when we modeled diffused exocytosis by allowing exocytosis to hit a larger number of mesh nodes, we did not observe a larger ring (Fig. S4d). Similar results also hold for simpler particle models (Fig. S5).

The SBE model (Fig. S4) and the particle models (Fig. S5) thus cannot explain the septin ring enlargement with diffused exocytosis, which we observed experimentally. Interestingly, when exocytosis is delayed via a conditional Bni1 allele, septin recruitment is reduced [Lai et al., 2018], suggesting that exocytosis aids in septin recruitment. To account for this, we expanded the SBE model to create the *Septin Binding and Exocytosis Recruitment* (SBER) model (Fig. 5a,k). To keep the SBER model simple, and as we do not know which protein might aid in vesicle-supported recruitment, we modeled that exocytosis delivered an unknown species (referred to as *X*), which recruits septin (Fig. 5a).

**Figure 5.**
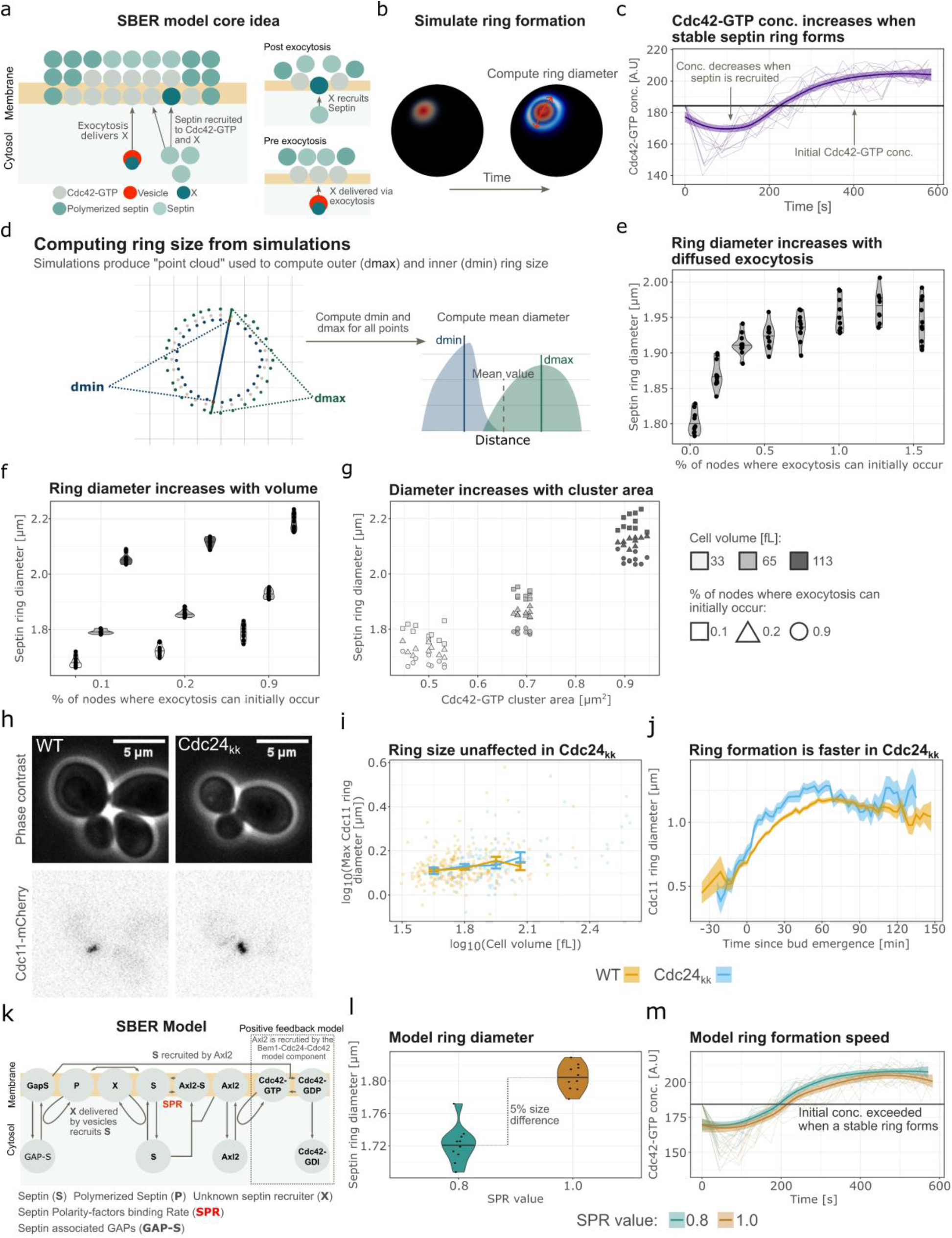
The *Septin Binding and Exocytosis Recruitment* (SBER) model explains how diffused exocytosis can produce a larger septin ring, and reveals the role of Cdc24 interaction with septins in septin ring formation. (a) Core concept of the SBER model. Septin represents the septin monomers (Cdc10, Cdc11, etc.) Briefly, septin is recruited towards the Cdc42-GTP cluster where it binds to polarity factors (see also for more details Fig. S4 and Fig. 5k), which prevents polymerization in the cluster center. This, combined with exocytosis displacing septin, promotes polymerization and ring formation at the cluster periphery. (b) A representative simulation example shows the Cdc42 polarization (red) and consecutive septin ring formation (blue). (c) Cdc42-GTP concentration for the SBER model plotted over time for n=10 simulations each initiated from the same Cdc42-GTP cluster with a loess-smoothing. (d) Schematic explanation of how septin ring diameter is computed from model simulations. (e) Septin ring diameter (*dmin+dmax)/2* for the SBER model plotted against the % of nodes that can initially be hit by exocytosis. The number of nodes that can be hit are those where the concentration of Cdc42 fulfils: *Cdc42-GTP > ε*max(Cdc42-GTP)*, where a smaller *ε* corresponds to more diffused exocytosis. For each condition, n=10 simulations, all starting from the same Cdc42-GTP cluster, were performed. In each case, the model was simulated for a long time to reach a stable ring, and then septin ring diameter was measured. (f) Septin ring diameter for the SBER model for three cell volumes and three different levels of diffused exocytosis. n=10 for each condition. (g) Septin ring diameter plotted against corresponding Cdc42-GTP cluster area for the simulations in panel (f). (h) Representative microscopy images of cells (phase contrast) and septin ring (Cdc11-mCherry) for WT and Cdc24_kk_ cells. (i) Cdc11 ring diameter measurements from microscopy images plotted against mother cell volume in double logarithmic scale for WT (n = 271 cells) and Cdc24_kk_ (n = 124 cells). Solid lines show binned means and error bars show standard error. (j) Mean Cdc11 ring diameter measurements from microscopy images plotted over the cell cycle for WT (n = 167 cells) and Cdc24_kk_ (n = 73 cells). Single cell traces are aligned at bud emergence. Ribbon shows standard error. (k) Schematic of the SBER model, where the Septin Polarity-factors binding Rate (SPR) is highlighted. The schematic focuses on the septin module, therefore, the Cdc42 polarity module is simplified on this schematic, for example, the Bem1-Cdc24-Cdc42-GTP complex that recruits Axl2 is omitted. (l) The final ring diameter is computed as in (d) for two different SPR values, at the final time-point in panel m (when the model has been simulated to a stable ring). For a larger SPR value, the ring diameter is approximately 5% larger. For each SPR value, n=10 simulations were performed. (m) Cdc42-GTP concentration over time for the same simulations as in panel (l). The thin lines correspond to individual simulations, and the thicker line to a loess smoothing.

Consistent with experimental observations [Okada et al., 2013], when simulating ring formation in the SBER model, the Cdc42-GTP concentration initially decreases upon septin recruitment and then increases after a stable ring has formed (Fig. 5c). In the Okada et al. model [Okada et al., 2013], such behavior was partially achieved by having septin create a diffusion barrier. However, the existence of such a barrier is debated [Sugiyama and Tanaka, 2019]. In the SBER model, a similar effect is obtained without a diffusion barrier, as the septin-recruited GAPs that associate with the septin ring convert any Cdc42-GTP that diffuses into the ring to Cdc42-GDP. This effectively concentrates Cdc42-GTPs inside the ring.

To investigate the effect of diffused exocytosis in our SBER model, we modeled exocytosis to be targeted towards the Cdc42 cluster, where vesicles could hit between 0.034% and 1.5% of the mesh nodes. Simulating 10 cells for each of eight conditions (Fig. 5b) and using our custom algorithm for ring diameter quantification (Fig. 5d), we found that the septin ring diameter gradually increases with more diffused exocytosis up to around 1% of nodes being hit (Fig. 5e). This suggests that stronger polarisome perturbations can lead to larger ring diameters.

Since the SBER model accounts for the increase in septin ring diameter observed in *bni1Δ* cells, we next tested whether it could also capture the observed increase of septin ring diameter with cell volume [Kukhtevich et al., 2020]. For three cell volumes (33, 65, 133 fL) and three levels of diffused exocytosis, we simulated 10 cells each. The septin ring diameter increased with both cell volume and the level of diffused exocytosis (Fig. 5f). Additionally, cluster area and ring diameter were positively correlated (Fig. 5g), in line with an increase in the Cdc42-cluster area driving the increase in the septin ring diameter.

Taken together, the SBER model shows that if exocytosis aids in septin recruitment, diffused exocytosis results in increased septin ring diameter. Furthermore, the SBER model recapitulates the experimentally observed increase of the septin ring diameter with the Cdc42-GTP cluster area and cell volume.

### Cdc24 interaction with septins is important for the timing of septin ring formation but not its final size

The SBER model suggests that diffused exocytosis accounts for the increased septin ring diameter observed in *bni1*Δ cells because exocytosis supports septin recruitment. We next asked what would be the effect of weakening the interaction between polarity proteins and septins. To this end, we further investigated with time-lapse imaging a Cdc24 mutant, Cdc24_kk_, which was reported to perturb the interaction between Cdc24 and septins [Chollet et al., 2020]. Both WT and Cdc24_kk_ mutant cells showed similar Cdc11 ring diameters and scaling of the ring diameter with cell volume (Fig. 5h, i). Next, we checked if there were any differences in the dynamics of septin ring formation. Interestingly, we found that in Cdc24_kk_ cells, the Cdc11 ring diameter reaches its maximum faster after bud emergence than in WT cells (Fig. 5j).

To better understand the mechanism underlying the faster formation of the septin ring, we turned again to computational modeling. By reducing the Septin Polarity-factors binding Rate (SPR), we weakened the interaction between septin and Axl2 in the SBER model (Fig. 5k), where, as mentioned above, Axl2 represents septin interacting polarity factors such as Cdc24. We then simulated 10 cells (Fig. 5l, m). Since in the model, the Cdc42-GTP concentration initially decreases and, after a stable ring is formed, increases (Fig. 5c), we decided to use the time it takes for Cdc42-GTP to exceed its initial concentration as a metric for ring formation speed. We found that consistent with our experimental observation, the Cdc42-GTP concentration increases faster for lower SPR values (Fig. 5m). This is likely because with a reduced SPR value, the inhibitory effect of polarity factors such as Cdc24 on septin polymerization is reduced, which promotes faster ring formation. Moreover, in the model, the septin ring diameter is also slightly smaller for lower SPR values (Fig. 5l). However, given the measurement error, such a small effect would be difficult to detect in experiments.

### Disruption of negative feedback leads to a larger Cdc42-GTP cluster area but not septin ring diameter

Earlier in this study, we showed that in Cdc24^38A^ cells (non-phosphorylatable negative feedback mutant), the Cdc42 cluster area is larger compared to WT cells (Fig. 2e, Fig. S3). We next addressed the role of the negative feedback in the regulation of the septin ring diameter. For this, we constructed new diploid strains in which one allele of Cdc10 was tagged with mCherry in WT and Cdc24^38A^ cells. Then we performed time-lapse experiments (Fig. 6a). Surprisingly, despite Cdc24^38A^ cells having a significantly larger Cdc42-GTP cluster, we did not find any major differences in the septin ring diameter at any point in the cell cycle (Fig. 6b, Fig. S6). Thus, we revealed a decoupling between the Cdc42-GTP cluster area and septin ring diameter.

**Figure 6.**
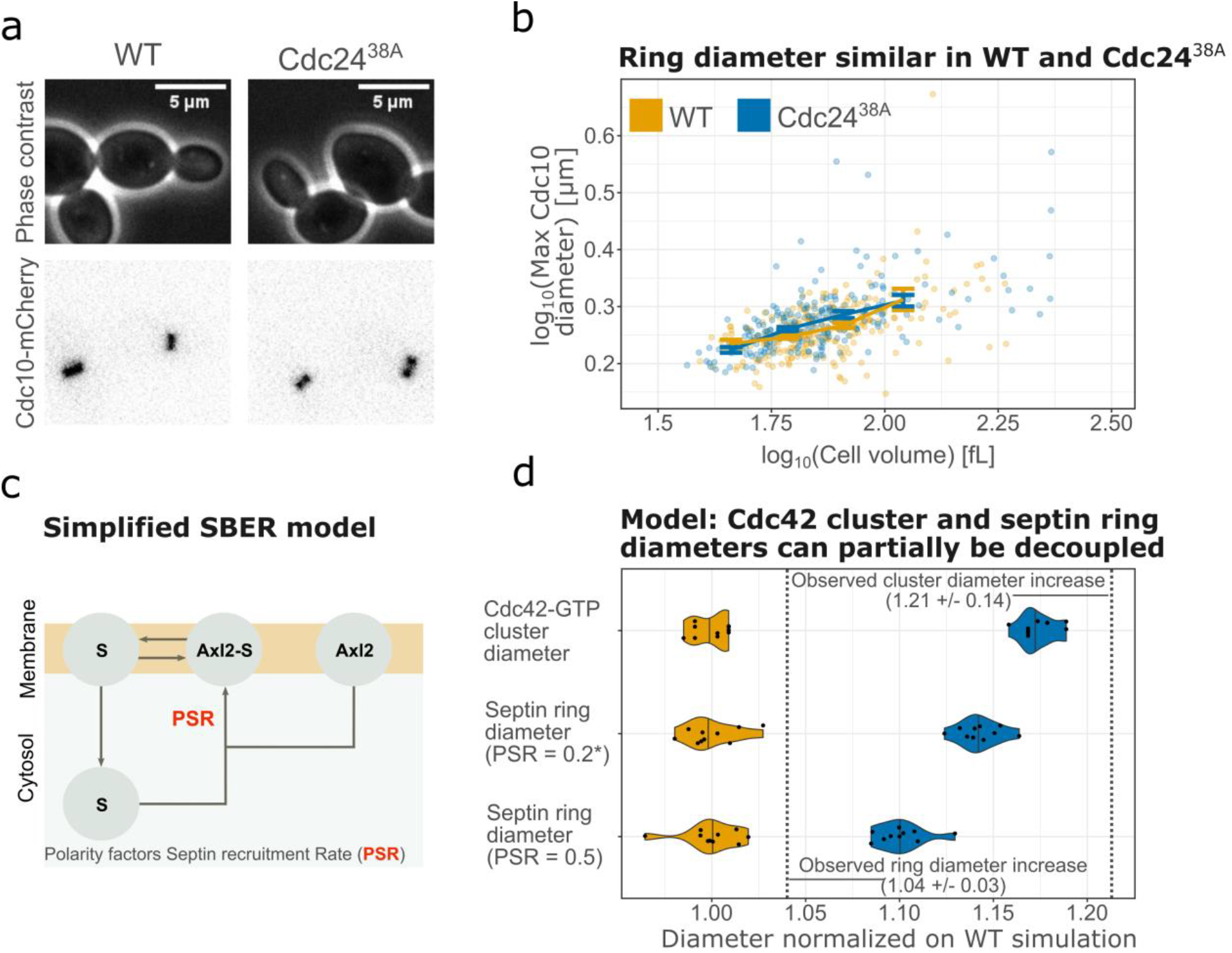
Septin ring diameter is decoupled from Cdc42-GTP cluster area in Cdc24^38A^ cells. (a) Representative microscopy images of cells (phase contrast) and septin ring (Cdc10-mCherry) for WT and Cdc24^38A^ cells. (b) Maximum Cdc10 ring diameter measurements from microscopy images plotted against corresponding mother cell volume in double logarithmic scale for WT (n = 237 cells) and Cdc24^38A^ (n = 240 cells). Solid lines show binned means and error bars show standard error. (c) Illustration of the PSR rate (Simplified version of Fig. 5k). (d) Cdc42-GTP cluster diameter and septin ring diameter for SBER model simulations of Cdc24^38A^ and WT, normalized on WT. Star denotes the default value (0.2) for the Polarity factors Septin recruitment Rate (PSR). The vertical dashed lines correspond to the increase in Cdc42 cluster and septin ring diameters observed experimentally (averaged across cell volume bins). For each condition, n=10 cells were simulated, and the model was simulated for a long time to reach a stable cluster area and septin ring.

To better understand this decoupling, we applied our SBER computational model. Previously, we found that reducing GAP activity in the positive feedback polarization model qualitatively captures the increased Cdc42-GTP cluster area observed in Cdc24^38A^ cells (Fig. S2). Moreover, given that we initiated SBER simulations from a Cdc42-GTP cluster, septin ring formation in the model is independent of the early polarization dynamics. Hence, to mimic Cdc24^38A^, we reduced GAP activity in the polarization module. We simulated 10 cells (Fig. 6c, d), and for ease of interpretation compared Cdc42-GTP cluster diameter, rather than area, with the septin ring diameter. The increase in ring diameter (fold change of 1.14 for PSR=0.2, Fig. 6d) was close to that in cluster diameter (1.17). By contrast, our experimental results showed that the septin ring diameter was almost intact (fold change of 1.04 for the maximum ring diameter), even though the maximum Cdc42-GTP cluster diameter (estimated as the square root of cluster area) increased 1.21-fold in Cdc24^38A^ averaged across cell volumes. To investigate if this discrepancy between experimental observations and the model could be explained by values chosen for the septin-related model parameters, we altered the activity of septin recruited GAPs, and septin concentration, but did not find any noticeable effect. However, when we increased the Polarity-factors Septin recruitment Rate (PSR parameter Fig. 6c), the increase in ring diameter compared to the previous simulations was smaller (fold change of 1.10 for PSR=0.5, Fig. 6d). This suggests that stronger recruitment of septins by polarity proteins can lead to a partial decoupling of Cdc42-GTP cluster diameter and ring diameter.

In summary, the Cdc42-GTP cluster area is increased in Cdc24^38A^ cells, while the Cdc10 ring diameter is largely the same. Thus, the Cdc24^38A^ mutation leads to a decoupling the Cdc42-GTP cluster area and the Cdc10 ring diameter.

## Discussion

The delicate interplay of positive and negative feedback enables yeast cells to cluster Cdc42-GTP at the presumptive bud site [Chiou et al., 2017], with an area that increases with cell volume [Kukhtevich et al., 2020]. Here, we showed that positive feedback, together with a volume-dependent increase in the amount of polarity proteins, is sufficient to explain this cluster scaling behavior (Fig. 1, 2a-d). Accordingly, in Cdc24^38A^ cells with disturbed negative feedback [Kuo et al., 2014], cluster area still scales with cell volume (Fig. 1g). Moreover, reanalysis of recently published proteomics data [Lanz et al., 2024] revealed that most polarity proteins have close to constant concentration with volume (Fig. 2d). Nevertheless, the exact mechanisms underlying the scaling of cluster area with cell volume in wild-type cells are not fully understood. In our study, we found that in both a model with only positive feedback and one with additional negative feedback that reduces Cdc42-GTP activation, Cdc42-GTP concentration in the cluster increases with cell volume. This could lead to a stronger membrane diffusion-driven flux that promotes cluster scaling with cell volume. Alternatively, in the negative feedback model when the negative feedback associated reaction rates were increased, the Cdc42-GTP concentration could plateau while the cluster area increases as the positive feedback recruits polarity proteins (Fig. 2e-h). Unraveling which scenario is at play requires quantification of the Cdc42-GTP concentration, but the commonly used approach for visualization of Cdc42-GTP species via tagged Gic2 does not provide sufficient accuracy to address this.

Yeast cells maintain the concentrations of polarity protein close to constant [Lanz et al., 2024] (Fig. 2d). This may serve the purpose of facilitating the formation of larger Cdc42-GTP clusters to achieve proper cell division as the nuclei increase in size with cell volume [Jorgensen et al., 2007]. Alternatively, and perhaps more likely, cluster scaling could be a side-effect of maintaining robust Cdc42 polarization. In our simulations with constant amounts of Cdc42 and the Bem1-complex, as well as constant GAP concentration, polarization fails (Fig. 2c). Overexpression of both Cdc42 and GEFs is fatal [Howel et al., 2012], and the deletion of GAP proteins reduces the replicative lifespan of yeast cells [Meitinger et al., 2014; Kang et al., 2022]. Even though Cdc42 polarization is remarkably robust [Brauns et al., 2023], there seems to be a need to maintain polarity protein balance. Interestingly, in computational models, an additional consequence of the increased amount of polarity proteins is that larger cells form multiple stable clusters [Borgqvist et al., 2021; Chiou et al., 2021], aligning with observations that in larger *rsr1*Δ cells (cells with random budding) multiple initial Cdc42 clusters form more frequently [Chiou et al., 2021]. With several initial clusters, bud site selection might become less efficient, which can also explain why in old, typically larger, cells bud site selection is less precise [Yang et al., 2022].

Following Cdc42-GTP cluster formation, septin ring formation initiates with septin recruitment to the bud site [Park and Bi, 2007]. Our prior work [Kukhtevich et al., 2020] revealed that deleting the formin and polarisome component Bni1 leads to an increased septin ring diameter despite an unchanged maximum Cdc42-GTP cluster area. Here, we further found that deleting other polarisome components, namely Spa2 or Bud6, causes larger ring diameters, while deleting the second formin Bnr1 showed no effect. Further, Bni1 had a larger effect than Spa2 and Bud6 (Fig. 3), possibly due to its direct interaction with F-actin [Liu et al., 2010; Xie et al., 2019]. We speculate that in *bnr1*Δ cells, Bni1 can compensate for Bnr1, consistent with previous observations [Yu et al., 2011; Garabedian et al., 2020]. An alternative explanation could be that formins can regulate the septin ring diameter only during a brief transient phase. Since Bnr1 is recruited later to the bud site [Pruyne et al., 2004], it may not be present at the bud site during this phase.

Given Bni1’s role as a formin involved in the assembly of F-actin cables [Park and Bi, 2007], and the suggestion by Okada et al. [Okada et al., 2013] that actin-dependent exocytosis regulates septin ring formation, we visualized F-actin and the exocyst subunit Exo84 [Jose et al., 2015] in both WT and *bni1Δ* cells. We found that the increased size of septin rings in *bni1Δ* cells is a possible consequence of exocytosis being diffused because of disturbed F-actin polarization toward the bud site (Fig. 4). Computational modeling showed that diffused exocytosis can produce a larger septin ring if exocytosis aids in septin recruitment (Fig. 5a-g), supporting previous observations that exocytosis might be involved in septin recruitment [Lai et al., 2018].

To account for the observation that septin ring formation can start without a clear exocytosis signal [Lai et al., 2018], our SBER model significantly extends the computational model by Okada et al. [Okada et al., 2013]. In addition to exocytosis promoting ring formation, and in line with the observation that Cdc42 inhibits septin polymerization *in vitro* [Sadian, 2013], in our model polarity proteins bind to septin and inhibit polymerization. This then drives septin to polymerize at the cluster periphery. A consequence of this second mechanism is that reducing the interaction between polarity factors and septin results in faster septin ring formation (Fig. 5k,m). Consistently, in a previously published Cdc24 mutant with a weakened interaction between Cdc24 and septins (Cdc24_kk_) [Chollet et al., 2020], we found that the septin ring diameter was unaffected, but that the time to reach the maximum diameter was shorter (Fig. 5i, j). Overall, given that the SBER model can explain several observed phenomena and that Cdc42 polarization is a remarkably robust process [Brauns et al., 2023], we argue that multiple mechanisms promote robust septin ring formation.

Lastly, we explored the effect on septin ring diameter when disrupting the negative Cdc42-GTP-centered feedback [Kuo et al., 2014] in Cdc24^38A^ cells. Surprisingly, we observed a decoupling between the Cdc42-GTP cluster area and the septin ring diameter, however in the opposite direction compared to *bni1Δ* cells [Kukhtevich et al., 2020]. In Cdc24^38A^ cells, the Cdc42-GTP cluster area increased but the septin ring diameter remained largely unaffected (Fig. 6). To understand why, we modified our SBER model to mimic Cdc24^38A^ cells by reducing GAP activity. In contrast to the experimental results, in the model, the septin ring diameter increased almost as strongly as the Cdc42-GTP cluster diameter (Fig. 6d). However, when we increased the recruitment rate of septin polarity factors in the SBER model, the septin ring diameter increased less than the Cdc42-GTP cluster diameter. In this case, stronger septin recruitment results in more septin-associated GAPs being recruited into the Cdc42-GTP cluster. Still, the decoupling predicted by the model was weaker than observed experimentally, highlighting that mechanisms beyond GAPs affect the interplay between septin and Cdc42. These mechanisms are likely less important if the negative feedback in the polarization pathway is present.

Taken together, by combining experiments and modeling, we identified key processes that contribute to setting the size of the budding yeast septin ring (Fig. 7). In particular, we found that positive feedback-based Cdc42-GTP cluster formation together with the amount of polarity proteins increasing with cell size is sufficient to explain the scaling of the septin ring diameter with cell volume. While this highlights the link between the Cdc42-GTP cluster and septin ring size, we also found that tuning specific parameters of the complex self-assembly process can lead to independent changes of both the Cdc42-GTP cluster area and septin ring diameter. Future studies will have to address whether such regulation is employed by cells to manipulate septin ring formation according to external or internal signals.

**Figure 7.**
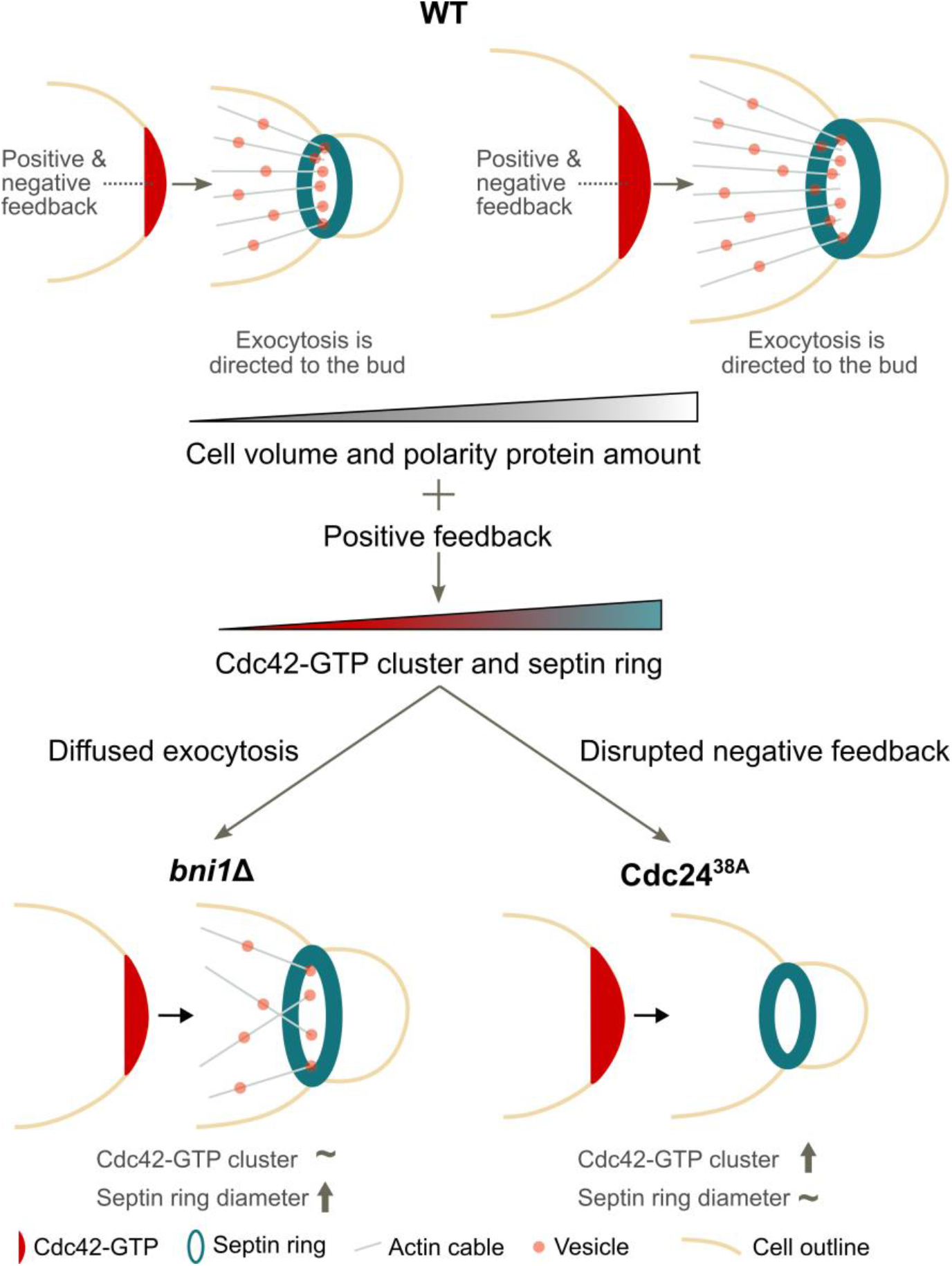
Illustration summarizing the mechanisms determining Cdc42-GTP cluster area and septin ring diameter. Positive feedback, together with polarity proteins increasing in amount as cell size increases, leads to an increase of Cdc42-GTP cluster area with cell volume. As a consequence, the septin ring diameter increases. The coupling of Cdc42-GTP cluster area and septin ring diameter can be disrupted by more diffused exocytosis or by a disrupted negative feedback in the Cdc42 polarization pathway.

## Methods

### Strain construction

*S. cerevisiae* strains were constructed using standard lithium acetate transformation. Detailed strain list can be found in Table S1. Where specified, we used *CDC10-mCITRINE, CDC10-mNEPTUNE2.5, CDC10-mCHERRY* or *CDC11-mCHERRY* to visualize the septin ring, *GIG2PDB-tdTOMATO* to visualize Cdc42-GTP, *ABP140-mCITRINE* to visualize F-actin, and *EXO84-mCITRINE* to visualize exocytosis. Strains with inducible Whi5 were used to tune cell size while maintaining cycling cell populations with similar doubling times [Kukhtevich et al., 2020; Claude et al., 2021]. Strains are available upon request.

### Growth conditions

Before experiments, cells were cultured in a synthetic complete liquid medium at 30 °C at low density to ensure exponential growth. As a carbon source, we used 2% glycerol and 1% ethanol (SCGE). After pre-culturing, strains carrying β-estradiol-inducible *WHI5* were grown in the presence of the respective β-estradiol concentration for ∼24 h before the start of the experiment, to ensure a steady state. Medium without β-estradiol was used during the live-cell microscopy experiments.

### Microfluidic devices

To acquire live single-cell data during time-lapse experiments, we used a previously reported custom-made microfluidic device that allows isolating cells in a dedicated region of interest and limits colony growth to the XY-plane [Kukhtevich et al., 2022]. The device includes eight separate cell culture chambers with a controllable medium exchange that enables parallel imaging of up to eight strains.

The microfluidic device was fabricated by means of standard soft lithography. Briefly, by using photolithography, a master mold for replication of the device design in polydimethylsiloxane (PDMS) was fabricated from SU-8 photoresist (MicroChem, USA) spin-coated on a 3″ Si wafer. The master mold was then filled with a 10:1 mixture of the base to curing agent of PDMS kit Sylgard 184 (Dow Corning, USA) and left at 60 °C for 4 h to crosslink the PDMS. After cross-linking, the PDMS replica was cut and peeled off from the master mold, and necessary inlets and outlets for tubing connections were made using a 1 mm puncher. Next, the replica was sealed with a coverslip after both were treated in O_2_ plasma.

### Live-cell microscopy

Live-cell time-lapse experiments (Figs. 1g, 2e, 3, 4a-g, 5h-j, 6a-b, S3, S6) were performed using the custom-made microfluidic device described above. Different strains were separately loaded in different chambers of the device. Constant medium flow at 20 μL/min was applied, enabling imaging of a colony growing over approximately 6 generations (10 h). Images were taken every 3 min for all time-lapse experiments, except for experiments imaging Abp140-mCitrine, for which a time interval of 10 min was used instead to allow for the acquisition of 5 z-slices with 1 μm steps. Temperature control was achieved by setting both a custom-made heatable insertion and an objective heater to 30 °C.

A Nikon Eclipse Ti-E with SPECTRA X light engine illumination and an Andor iXon Ultra 888 camera were used for epifluorescence microscopy. A plan-apo λ 100x/1.45NA Ph3 oil immersion objective was used to take phase contrast and fluorescence images. mCitrine fluorescence was imaged by exposure for 400 ms, illuminating with the SPECTRA X light engine at 504 nm and about 12 mW (20%) power for all experiments, except for experiments imaging Abp140-mCitrine, for which the exposure time was set to 300 ms and the power to about 24 mW (40%). tdTomato fluorescence was imaged by exposure for 200 ms, illuminating with the SPECTRA X light engine at 556 nm and about 26 mW (10%) power. mCherry fluorescence was imaged by exposure for 200 ms, illuminating with the SPECTRA X light engine at 556 nm and about 26 mW (10%) power. mNeptune2.5 fluorescence was imaged by exposure for 400 ms, illuminating with the SPECTRA X light engine at 556 nm and about 26 mW (10%) power for all experiments except for experiments imaging Abp140-mCitrine for which the exposure time was set to 300 ms. Fluorescence of all fluorescent proteins was detected using suitable emission wavelength filters.

### Quantification and statistical analysis

#### Cell segmentation and tracking

For experiments shown in Figs. 1, 2e, 4d-g, 5h-j, 6, S3, S6 cell segmentation, cell volume calculations, lineage annotations and cell-cycle stage assignments were performed using the Cell-ACDC software available at https://github.com/SchmollerLab/Cell_ACDC [Padovani et al., 2022]. More specifically, we used the YeaZ [Dietler et al., 2020] neural network option in Cell-ACDC for segmentation and tracking and manually corrected where necessary.

For experiments shown in Fig. 4a-c, cells were automatically segmented and tracked based on phase-contrast images using the Matlab-based Phylocell software [Fehrmann et al., 2013]. The results of automatic segmentation and tracking were visually inspected and manually corrected if necessary.

For experiments shown in Fig. 3, cells were segmented and tracked based on phase-contrast images using the Matlab-based software described in [Doncic et al., 2013]. The result was manually checked for each cell included in the analysis, and poorly segmented or wrongly tracked cells were rejected from the analysis.

#### Calculation of cell volume and length along the major axis

Cell volume and length along the major axis was calculated based on 2D phase contrast images as described previously [Kukhtevich et al., 2020, Padovani et al., 2022]. Briefly, cell contours were aligned along their major axis, and divided into slices perpendicular to the major axis. To estimate cell volume, we then assumed rotational symmetry of each slice around its middle axis parallel to the cell’s major axis.

#### Analysis of the septin ring diameter and Exo84 cluster diameter

For the experiments shown in Fig. 3, an analysis of the septin ring diameter was performed using Fiji and Matlab, as previously described [Kukhtevich et al., 2020]. Briefly, Fiji was used to automatically determine the position and orientation of the ring and obtain an intensity line profile along its major axis. This intensity profile was then further analyzed using Matlab to quantify the ring diameter.

For the experiments shown in Fig. 4a-c, the septin ring diameter based on Cdc10-mNeptune2.5 fluorescence was calculated from the five frames centered around the time of clear bud emergence, using an approach previously described in [Kukhtevich et al., 2020]. To do so, a fluorescence profile along a cell segmentation contour was taken for each frame after applying the 3 × 3 mean filter on the fluorescence image. A mean fluorescence profile was then calculated based on the five selected frames. Finally, we used the full width at half maximum as an estimate for the septin ring diameter. For this, the minimum signal in the fluorescence profile was defined as the baseline. All length profiles were visually inspected and cells were rejected from further analysis when this approach resulted in obvious artifacts.

The septin ring diameters shown in Figs. 4f-g, 5i, 5j, 6b, S6 and the Exo84 cluster diameter shown in Fig. 4e,g were measured as follows: first, the cells were segmented, tracked over time, and their cell cycle progression was annotated from phase contrast signal using the software Cell-ACDC [Padovani et al., 2022]. Specifically, for segmentation and tracking we used the model YeaZ (embedded in Cell-ACDC) [Dietler et al., 2020]. Segmentation and tracking were visually inspected, and errors were corrected with Cell-ACDC. The cell cycle progression was determined by annotating bud emergence, mother-bud pairs, and cell division. Note that cell division is visually determined by carefully checking for sudden bud movement (indicating the bud divided from the mother cell). Next, we used an automatic custom Python routine to calculate the ring/cluster diameter for each complete cell cycle. In the first step, the algorithm applies a Gaussian filter (with ‘sigma = 2.0‘) to each frame of the video. Next, the routine extracts the intensities from an elliptical region whose longer axis is aligned with the line connecting the centers of mass of the mother and bud, while the center lies on the contact point between the mother and bud. The contact point is determined as the point along the line connecting the centers of mass where the segmentation ID changes. The shorter axis length was set to 10 pixels, while the longer axis of the ellipse was calculated as the distance between the two points intersecting the long axis and the contour of the hull image of the mother-bud object. These two distances were selected to ensure that the brightest part of the ring/cluster is included in the elliptical area. In the second step, the intensities from this area are sorted in descending order, and the 10th value from maximum intensity is taken as a representative data point for each timeframe to build a cell cycle curve over time. Note that other values were tested, including the mean, the max, the median, the 20th value from maximum, etc. and we found the 10th value to be the more robust for the next step. In the third step, a threshold value was determined for each cell cycle as the mean between maximum and minimum in the curve. In the fourth step, the ring/cluster structure was segmented for each time point and cell by applying a threshold on the Gaussian filtered intensities in each mother-bud object using the threshold value determined in the previous step. In the fifth step, for each timepoint, only the cells where one (for Exo84 signal) or a maximum of two (for septin ring signal) objects overlapping by at least 50% with the elliptical region (see above) were kept for the analysis. Note that the overlap was calculated as the intersection-over-area ratio where intersection is the number of pixels in both the elliptical region and the sub-cellular objects, while area is the number of pixels in the sub-cellular objects mask.

Finally, the ring diameter/Exo84 cluster diameter is determined as the major axis length of the segmented object using the scikit-image function `skimage.measure.regionprops` [van der Walt et al., 2014]. The maximum of the septin ring/Exo84 cluster diameter was determined from the evolution of the diameter over time. Cells whose S/G2/M phase was not fully tracked and the tracked duration was less than 45 minutes, and cells for which the ring/cluster segmentation was not successful for at least 3 or 2 frames, respectively, were discarded from the analysis. For the ring diameter over time plots in Figs. 5j and S6a, cells without fully tracked G1 phase were discarded.

#### Analysis of the Cdc42

To determine the Cdc42 cluster areas in Figs. 1g, 2e, and S6b, we developed an automatic custom Python routine that starts from fully annotated cell cycles of single cells and is based on our previously published method [Kukhtevich et al., 2020]. See the previous section for more details about how the cell pedigrees were annotated. In the first step, the algorithm applies a gaussian filter (with ‘sigma = 2.0‘) to each frame of the video. Next, the routine determines a threshold value for each cell using the following formula:

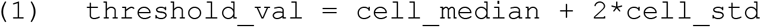

where `cell_median` and `cell_std` are the median and the standard deviation of the gaussian filtered intensities in each cell, respectively. In the second step, the algorithm thresholds the intensities from each cell using the threshold values determined in the previous step to achieve segmentation of the clusters. In the third step, the algorithm identifies individual segmented objects in the cell and keeps only the largest one. Finally, comparing the different time points, the routine extracts the maximum cluster area prior to bud emergence from cells with fully tracked G1 phase. The intensity of the brightest pixel in the cluster was used as a measure for the maximum Cdc42-GTP intensity in Figs. 2e and S3b. For the Cdc42-GTP cluster area over time plot in Fig S6b, cells without fully tracked G1 were discarded from the analysis.

#### Analysis of the Abp140 cluster

All parameters of the Abp140 cluster (Fig. 4b, c) were calculated for the three frames centered around the frame with the peak Abp140 localization, which was determined by visual inspection.

To measure the Abp140 cluster area, we applied a thresholding approach similar to that used by Okada et al. to measure Cdc42-GTP cluster area [Okada et al., 2013, Kukhtevich et al., 2020]. For each cell and each time point, a threshold was defined as the median value + 2 standard deviations of the fluorescence pixel intensities within the selected cell. All pixels with a value higher than this threshold were then counted as part of the cluster, and the cluster area was defined as the median number of pixels in the cluster across the three analyzed frames.

### Computational modeling

#### Positive feedback model

The positive feedback model consists of the reactions in Tab. 1. It was originally published in [Woods et al., 2015], and is based on the model of Goryachev and Pokhilko [Goryachev and Pokhilko, 2008]. Model species are Cdc42-GTP (*Cdc42T*, membrane-bound), Cdc42-GDP (*Cdc42D*, membrane-bound), Cdc42-GDI (*Cdc42c*, cytosolic), Bem-GEF-Cdc42-GTP complex (*BemGEF42*, membrane-bound), and the Bem-GEF/Cdc24 complex (*BemGEFm* membrane-bound and *BemGEFc* cytosolic).

**Table 1.** Model reactions and kinetic parameter values positive feedback model.

| Reaction | Parameter | Value | Reference |
| --- | --- | --- | --- |
| $BemGEFc \rightarrow BemGEFm$ | $k_{1a}$ | 10 | [Chiou et al., 2021] |
| $BemGEFm \rightarrow BemGEFc$ | $k_{1b}$ | 10 | [Chiou et al., 2021] |
| $Cdc42D + BemGEFm \rightarrow Cdc42T + BemGEFm$ | $k_{2a}$ | 0.16 | [Chiou et al., 2021] |
| $Cdc42T \rightarrow Cdc42D$ | $k_{2b}$ | 0.35 | [Chiou et al., 2021] |
| $Cdc42D + BemGEF42 \rightarrow Cdc42T + BemGEF42$ | $k_3$ | 0.35 | [Chiou et al., 2021] |
| $BemGEF + Cdc42T \rightarrow BemGEF42$ | $k_{4a}$ | 10 | [Chiou et al., 2021] |
| $BemGEF42 \rightarrow BemGEFm + Cdc42T$ | $k_{4b}$ | 10 | [Chiou et al., 2021] |
| $Cdc42I \rightarrow Cdc42D$ | $k_{5a}$ | 36 | [Chiou et al., 2021] |
| $Cdc42D \rightarrow Cdc42I$ | $k_{5b}$ | 0.65 | [Chiou et al., 2021] |
| $BemGEFc + Cdc42T \rightarrow BemGEF42$ | $k_7$ | 10 | [Chiou et al., 2021] |

Assuming much faster cytosolic than membrane diffusion (*D*_*c*_ → ∞), the reactions in Tab. 1 comprise a coupled system of ordinary differential equations (ODEs) and partial differential equations (PDEs):

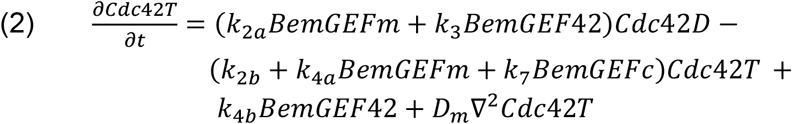

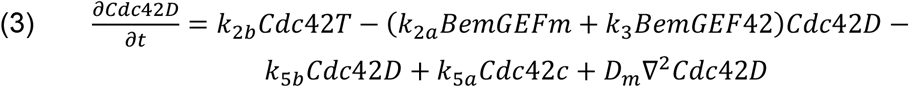

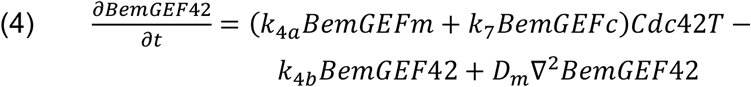

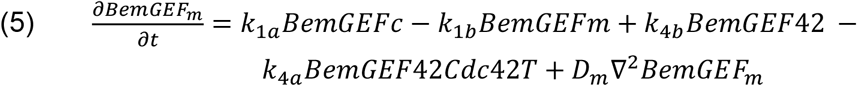

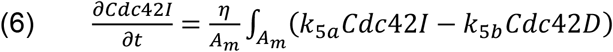

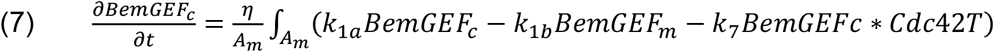

where *A*_*m*_ is the cell-surface area, and *η* is the membrane-to-cytoplasm volume ratio *V*_*m*_/*V*_*c*_. We assume a membrane thickness of *R*_*m*_ ≈ 10 nm [Goryachev and Pokhilko, 2008] and thereby *η* can be computed as:

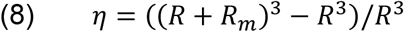

Thus, the membrane is treated as a compartment and the concentration for surface species refers to the concentration in the membrane compartment.

#### Negative feedback model

The positive feedback model can be expanded to include negative feedback. The resulting negative feedback model consists of the reactions in Tab. 2, and was originally published in [Kuo et al., 2014]. The model species are Cdc42-GTP (*Cdc42T*, membrane-bound), Cdc42-GDP (*Cdc42D*, membrane-bound), Cdc42-GDI (*Cdc42c*, cytosolic), Bem-GEF-Cdc42-GTP complex (*BemGEF42*, membrane-bound), and the Bem-GEF/Cdc24 complex (*BemGEFm* membrane-bound and *BemGEFc* cytosolic). Asterisks, e.g. *BemGEF42\**, denote phosphorylated species.

**Table 2.** Model reactions and kinetic parameter values negative feedback model.

| Reaction | Parameter | Value | Reference |
| --- | --- | --- | --- |
| $BemGEFc \rightarrow BemGEFm$ | $k_{1a}$ | 10 | [Chiou et al., 2021] |
| $BemGEFm \rightarrow BemGEFc$ | $k_{1b}$ | 10 | [Chiou et al., 2021] |
| $Cdc42D + BemGEFm \rightarrow Cdc42T + BemGEFm$ | $k_{2a}$ | 0.16 | [Chiou et al., 2021] |
| $Cdc42T \rightarrow Cdc42D$ | $k_{2b}$ | 0.35 | [Chiou et al., 2021] |
| $Cdc42D + BemGEF42 \rightarrow Cdc42T + BemGEF42$ | $k_3$ | 0.35 | [Chiou et al., 2021] |
| $BemGEF + Cdc42T \rightarrow BemGEF42$ | $k_{4a}$ | 10 | [Chiou et al., 2021] |
| $BemGEF42 \rightarrow BemGEFm + Cdc42T$ | $k_{4b}$ | 10 | [Chiou et al., 2021] |
| $Cdc42I \rightarrow Cdc42D$ | $k_{5a}$ | 36 | [Chiou et al., 2021] |
| $Cdc42D \rightarrow Cdc42I$ | $k_{5b}$ | 0.65 | [Chiou et al., 2021] |
| $BemGEFc + Cdc42T \rightarrow BemGEF42$ | $k_7$ | 10 | [Chiou et al., 2021] |
| $BemGEF^* \rightarrow BemGEFc^*$ | $k_{1b}$ | 10 | [Chiou et al., 2021] |
| $BemGEF^* + Cdc42T \rightarrow BemGEF42^*$ | $k_{4a}$ | 10 | [Chiou et al., 2021] |
| $BemGEF42^* \rightarrow BemGEF^* + Cdc42T$ | $k_{4b}$ | 10 | [Chiou et al., 2021] |
| $BemGEF42 + BemGEF42 \rightarrow BemGEF42 + BemGEF42^*$ | $k_{8max}, k_{8h}$<br>$k_{8n}$ | 0.0063,6<br>10 | [Chiou et al., 2021] |
| $BemGEF42 + BemGEF42^* \rightarrow BemGEF42^* + BemGEF42^*$ | $k_{8max}, k_{8h}$<br>$k_{8n}$ | 0.0063,6<br>10 | [Chiou et al., 2021] |
| $BemGEFc^* \rightarrow BemGEFc$ | $k_{9max}, k_{9h}$<br>$k_{9n}$ | 0.0044,6,<br>10.003 | [Chiou et al., 2021] |

The difference of the negative feedback model compared to the positive feedback model is the presence of one delayed negative feedback, which via a Hill activation is triggered when *BemGEF42* activity is sufficiently high. Briefly, when *BemGEF42* activity gets too high, it autophosphorylates, and in the phosphorylated state, it cannot promote *Cdc42T* production. To become dephosphorylated, *BemGEF\** must be recycled into the cytosol. That dephosphorylation only occurs in the cytosol creates a delayed negative feedback, which besides reducing Cdc42-GTP activation facilitates, as observed experimentally [Howell et al., 2012], oscillations in Cdc42-GTP cluster intensity during polarization. Overall, the model equations are:

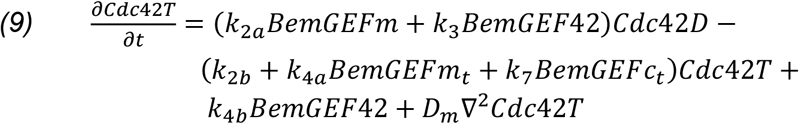

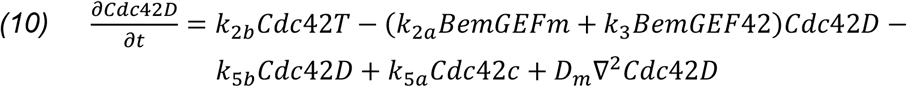

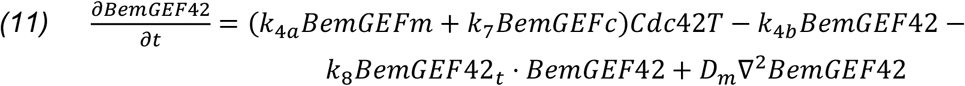

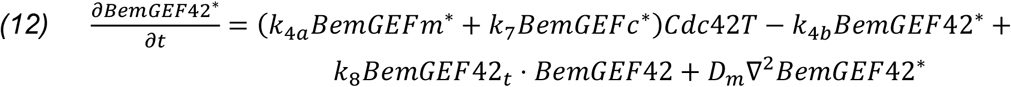

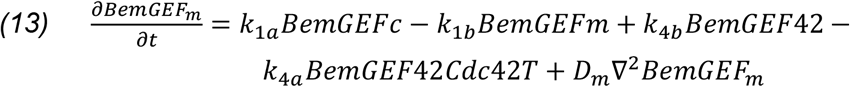

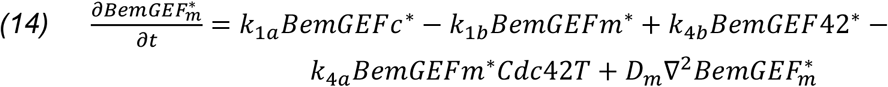

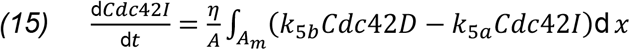

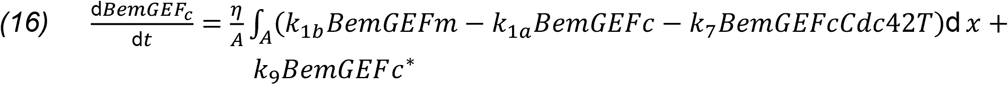

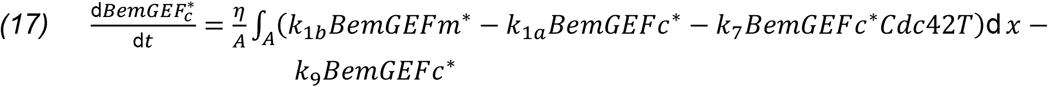

#### Alternative positive feedback model

The alternative Cdc42 positive feedback model (Fig. S1a) is based on the reactions in Tab. 3 and was published in [Borgqvist et al., 2021]. The model includes three species: Cdc42-GTP (*Cdc42T*, membrane-bound), Cdc42-GDP (*Cdc42D*, membrane-bound), and Cdc42-GDI (*Cdc42I*, cytosolic). The third-order reaction (reaction 5 in Tab. 3), is a simplification of the positive feedback mechanisms into a single reaction step. The parameters for this model were derived in a manuscript under preparation.

**Table 3.** Model reactions for alternative positive feedback model.

| Reaction | Parameter | Value | Reference |
| --- | --- | --- | --- |
| $Cdc42I \rightarrow Cdc42D$ | $k_1, k_{max}$ | 0.28, 2.54 | MS in preparation |
| $Cdc42D \rightarrow Cdc42I$ | $k_{-1}$ | 0.133 | MS in preparation |
| $Cdc42D \rightarrow Cdc42T$ | $k_2$ | 0.001368 | MS in preparation |
| $Cdc42T \rightarrow Cdc42D$ | $k_{-2}$ | 0.028 | MS in preparation |
| $2Cdc42T + Cdc42D \rightarrow 3Cdc42T$ | $k_3$ | 1 | MS in preparation |
| Membrane diffusion<br>*Cdc42T* | $D_T$ | 0.011 | MS in preparation |
| Membrane diffusion<br>*Cdc42D* | $D_D$ | 1.21 | MS in preparation |
| Cytosolic diffusion<br>*Cdc42I* | $D_I$ | 10 | MS in preparation |

If reactions are turned into equations, we obtain:

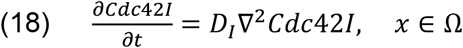

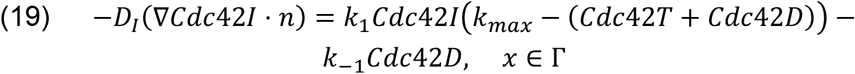

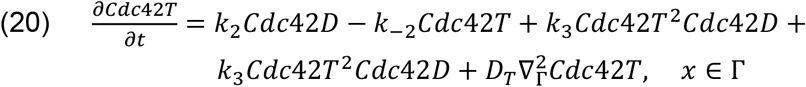

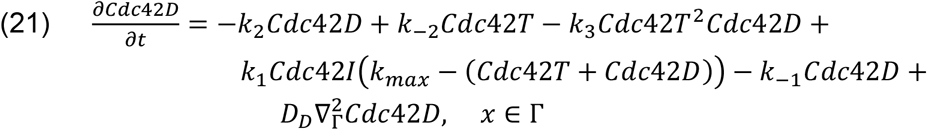

Here Ω refers to the cytosol, and Γ to the membrane. Note that compared to other models, cytosolic diffusion is here treated as finite, and the cytosol is part of the model geometry (Fig. S1).

#### Septin ring models

The septin ring model consists of the reactions in Tab. 4. Model species are Cdc42-GTP (*Cdc42T*, membrane-bound), Cdc42-GDP (*Cdc42D*, membrane-bound), Cdc42-GDI (*Cdc42c*, cytosolic), Bem-GEF/Cdc24 complex (*BemGEFm* membrane-bound and *BemGEFc* cytosolic), Bem-GEF-Cdc42-GTP complex (*BemGEF42*, membrane-bound), Axl2 (*Axl2* membrane-bound and *Axl2c* cytosolic), septin-associated Gap (*GapS* membrane-bound and *GapSc* cytosolic), monomeric septin (*S* membrane-bound and *Sc* cytosolic), polymerized septin (*P*, membrane-bound), and septin recruiter (*X*, membrane-bound). For a motivation of the model structure see supplementary information.

**Table 4.** Model reactions for the SBE and SBER models. Note, the SBE model is identical to the SBER model, except it does not include any of the reactions with species *X*.

| Reaction | Parameter | Value | Reference |
| --- | --- | --- | --- |
| $BemGEFc \rightarrow BemGEFm$ | $k_{1a}$ | 10 | [Chiou et al., 2021] |
| $BemGEFm \rightarrow BemGEFc$ | $k_{1b}$ | 10 | [Chiou et al., 2021] |
| $Cdc42D + BemGEFm \rightarrow Cdc42T + BemGEFm$ | $k_{2a}$ | 0.16 | [Chiou et al., 2021] |
| $Cdc42T \rightarrow Cdc42D$ | $k_{2b}$ | 0.63 | [Ghose et al., 2021] |
| $Cdc42D + BemGEF42 \rightarrow Cdc42T + BemGEF42$ | $k_3$ | 0.35 | [Chiou et al., 2021] |
| $BemGEF + Cdc42T \rightarrow BemGEF42$ | $k_{4a}$ | 10 | [Chiou et al., 2021] |
| $BemGEF42 \rightarrow BemGEFm + Cdc42T$ | $k_{4b}$ | 10 | [Chiou et al., 2021] |
| $Cdc42I \rightarrow Cdc42D$ | $k_{5a}$ | 144 | [Ghose et al., 2021] |
| $Cdc42D \rightarrow Cdc42I$ | $k_{5b}$ | 20.8 | [Ghose et al., 2021] |
| $BemGEFc + Cdc42T \rightarrow BemGEF42$ | $k_7$ | 10 | [Chiou et al., 2021] |
| $GapSc + P \rightarrow GapS + P$ | $k_{12a}$ | 10 | [Chiou et al., 2021] |
| $GapS \rightarrow GapSc$ | $k_{12b}$ | 10 | This work |
| $GapS + Cdc42T \rightarrow GapS + Cdc42D$ | $k_{13}$ | 1.5 | This work |
| $Axl2S \rightarrow Axl2 + S$ | $k_{15}$ | 1.0 | This work |
| $S + S \rightarrow 2P$ | $k_{16}$ | 0.05 | This work |
| $S + P \rightarrow P + P$ | $k_{17}$ | 0.125 | This work |
| $P \rightarrow S$ | $k_{18}$ | 0.1 | This work |
| $Axl2 + S \rightarrow Axl2$ | $k_{19}$ | 4.5 | This work |
| $Axl2 + Sc \rightarrow Axl2S$ | $k_{20}$ | 0.2 | This work |
| $S \rightarrow Sc$ | $k_{21}$ | 0.65 | This work |
| $Axl2 \rightarrow Axl2c$ | $k_{22}$ | 10.5 | This work |
| $BemGEF42 + Axl2c \rightarrow BemGEF42 + Axl2$ | $k_{23}$ | 26 | This work |
| $X \rightarrow \phi$ | $k_{24}$ | 0.1 | This work |
| $X + Sc \rightarrow X + Sc$ | $k_{25}$ | 5.5 | This work |
| Membrane diffusion | $D_m$ | 0.0025 | [Chiou et al., 2021] |
| Membrane diffusion P and X | $D_p$ | 0.00025 | This work |

Model parameters were tuned with the following rationale:

- Axl2 recruitment parameters (*k*_22_, *k*_23_) were set to allow relatively rapid recycling between membrane and cytosol, to mimic the behavior of other polarity components such as Cdc42.
- Axl2 septin recruiting parameters (*k*_15_, *k*_19_, *k*_20_) were set to enable sufficiently fast septin recruitment. For example, if *k*_20_ is too small, ring formation does not take off. If *k*_15_ is too large, septin polymerizes too strongly in the pole center, causing the Cdc42-GTP cluster to collapse due to the septin recruited GAP proteins.
  - *k*_19_ is in the manuscript also referred to as the Septin Polarity-factors binding rate (SPR)
  - *k*_20_ is in the manuscript also referred to as the Polarity-factors septin recruitment Rate (PSR).
- Septin polymerization parameters (*k*_18_, *k*_19_) were selected to allow sufficiently fast polymerization. These, along with Axl2 septin binding parameters (*k*_15_, *k*_19_, *k*_20_), were kept small enough to prevent strong polymerization in the Cdc42 cluster center which causes cluster collapse, but large enough to allow ring formation.
- Septin-associated Gap (GAP-S) parameters (*k*_12*a*_, *k*_12*b*_, *k*_13_) were chosen to facilitate sufficiently fast recruitment of GAP-S. Parameter *k*_13_ was set strong enough to allow the septin ring to “capture” Cdc42-GTP, acting as a substitute for a diffusion barrier (see Fig. 5k).
- *X*-associated parameters (*k*_24_, *k*_25_) were adjusted to ensure a wider pole of *X* when exocytosis is diffused (see supplementary).
- Parameters *k*_2*b*_, *k*_5*a*_ *and k*_5*b*_ differ from the model used for assessing pole size (Tab. 1). This adjustment was made to make the pole more robust to exocytosis, and to obtain a more realistic smaller cluster (important for forming a septin ring in a realistic context). For assessing pole size, we aimed to maintain similar parameters to the negative feedback model for comparison.

Overall, if the model reactions are translated into equations we get:

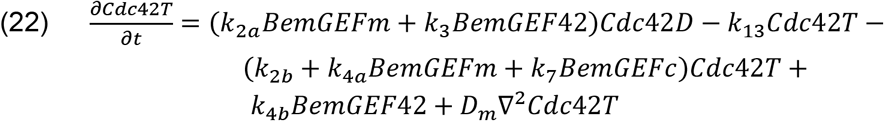

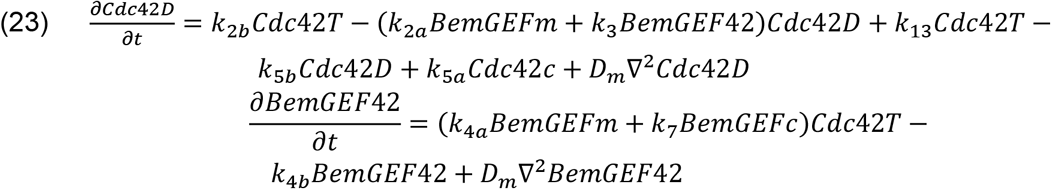

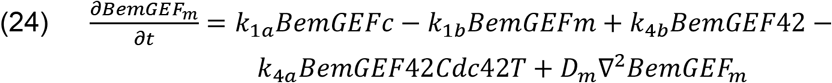

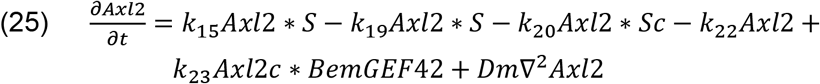

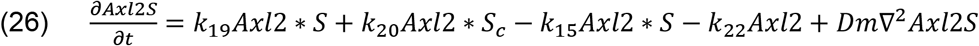

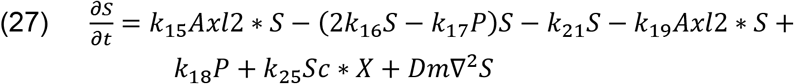

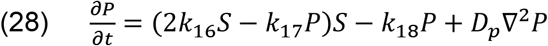

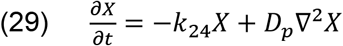

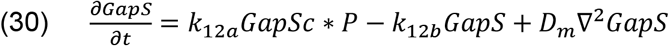

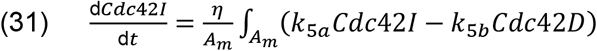

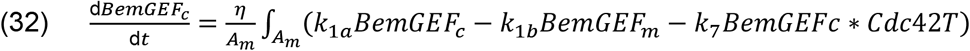

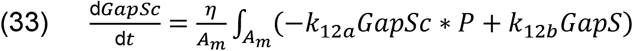

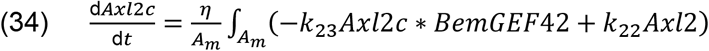

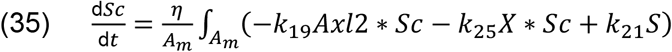

In addition to chemical reactions, the model includes exocytosis. Following experimental observations [Watson et al., 2014; Ghose et al., 2021], vesicles are modeled to deliver Cdc42-GDP at a lower concentration than the current Cdc42 total concentration in the pole (100 μM compared to around 180 μM). Thus, they effectively dilute the pole. *X* is delivered at a concentration of 20 μM. The nodes that can be hit by exocytosis are those where the concentration of Cdc42 fulfills: *Cdc42-GTP > ε*max(Cdc42-GTP)*, where a smaller *ε* corresponds to more diffuse exocytosis, and for each exocytosis event one of these nodes is randomly selected.

#### Exocytosis modeling

Exocytosis is modeled using the approach described in [Gerganova et al., 2021]. Here, exocytosis displaces proteins radially away from the center of exocytosis. The displacement for a molecule at an arc length (geodesic distance) *s* from the center of exocytosis is given by:

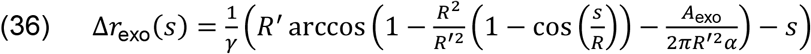

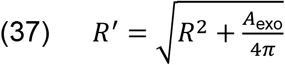

where *R’* is the radius of the sphere following exocytosis, *R* is the radius of the cell, *A*_*exo*_ is the vesicle surface area, *γ* and *α* are hydrodynamic parameters. Specifically, *α* represents the fraction of the fluid component in the membrane and *γ* ≤ 1 is the ratio of lipid to protein velocity. Like in previous studies on *S. pombe*, we set *γ* = 1 and *α=0.5* [Gerganova et al., 2021]. Additionally, following earlier work in *S. cerevisiae*, we use *r*_*exo*_ = 50 nm [Ghose et al., 2021]. A summary of parameter values can be found in Tab. 5.

Exocytosis recruitment is modeled as a stochastic event with a rate *λ*. Previous modeling on septin ring formation used a rate of 0.2/s [Okada et al., 2013]. Measurements under no growth in *S. pombe* yielded a rate of 0.5/s, and for *S. cerevisiae* during yeast polarization the rate was measured to be around 0.41/s [Layton et al., 2011], while when modeling polarity patch movement a rate of 0.83/s has been used [Ghose et al., 2021]. Considering that most estimates are larger than the value in [Okada et al., 2013], we opted for a compromise and set the rate to 0.4/s.

Applying the same principle, we can extend the model to incorporate endocytosis. However, we exclude endocytosis due to three reasons. Firstly, due to the smaller size of endosomes (approximately 1/4 the area of exosomes, resulting in a comparatively smaller impact from a single endocytosis event). Secondly, due to their wider occurrence area (thus exerting less influence on ring formation and the pole) [Layton et al., 2011]. Thirdly, due to the uncertainty regarding whether they recycle any noticeable polarity proteins (which could affect polarization). Overall, as in previous septin ring modeling [Okada et al., 2013], we made the choice to exclude endocytosis, which further reduces simulation time.

**Table 5.** Exocytosis model parameters for the SBE and SBER models.

| Parameter | Value | Interpretation | Reference |
| --- | --- | --- | --- |
| $\alpha$ | 0.5 | Mobile membrane fraction | [Gerganova et al., 2021] |
| $\gamma$ | 1.0 | Ratio of lipid to protein velocity | [Gerganova et al., 2021] |
| $r_{exo}$ | 50nm | Exosome radius | [Layton et al., 2011] |
| $\lambda$ | 0.4/s | Exocytosis rate | This work, based on; [Layton et al., 2011; Okada et al., 2013; Gerganova et al., 2021]. |

#### Simple particle model

In the simple particle model (Fig. S5a), we initialize the simulation with 1000 particles randomly distributed within a pole that occupies a fraction *Ω* of the cell-surface area. Subsequently, we run the simulation for 20 iterations, applying exocytosis following the approach described above. Different values for the tuning parameters *α* and *γ* (Tab. 5) were explored, and the tested parameter values are provided in the first three rows of Tab. 6.

Exocytosis is permitted within a cluster, denoted as Σ, which occupies a fraction *Ω* of the cell area (the same pole where particles start). To simulate varying degrees of diffused exocytosis, we initially generate 10,000 random points within Σ. The exocytosis site is then randomly sampled from these points, with each point weighted by:

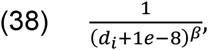

where *d*_*i*_ represents the distance of point *i* to the center of Σ. For small values of β, exocytosis is widespread, while larger values make it more concentrated (Fig. S5b). Alternatively, the exocytosis hit site could be modeled using a Gaussian distribution, as done on the plane in [Ghose et al., 2021]. However, deriving the Gaussian distribution on the surface of a sphere (as in our simulations) is challenging, therefore we opted for the approach above.

#### More complex particle model

In the more complex particle model (Fig. S5d), particles are randomly recruited within a cluster denoted as Σ, which occupies a fraction *Ω* of the cell area. Particles are recruited at the rate *k*_*rec*_, they diffuse on the surface with the rate *D*_*m*_, and disassociate with a rate *k*_*off*_. Exocytosis is modeled using the same approach as in the simple particle simulator.

The simulation algorithm is custom-made. Specifically, recruitment is simulated using τ-leaping [Gillespie, 2007] with a step length of *dt=0.01 s*. Recruited particles are randomly distributed within Σ. Diffusion is modeled with the same time step; *dt*. Once a particle is recruited to the membrane, its membrane lifetime *t*_*i*_ is determined using the SSA (Gillespie) algorithm [Gillespie, 2007], as the particles are modeled as independent (i.e., they do not interact). Once the simulation time *t* exceeds *t*_*i*_, the particle is removed from the sphere surface.

Despite having fewer species and reactions than the SBER model (Fig. 5a, k), the complex particle model captures key properties of septin ring formation. Particles are recruited within a defined area (denoted as Σ, mimicking the Cdc42-GTP cluster). They undergo diffusion, dissociate from the membrane, and experience displacement via exocytosis. For the parameters used see Tab. 6.

**Table 6.** Tested values for particle simulator. The values column corresponds to tested values in simulations. Note for Ω only results for 0.03 are presented, as results were consistent for different values.

|  | Values | Interpretation |
| --- | --- | --- |
| $\beta$ | 0.1 - 3.0 | Parameter deciding how concentrated exocytosis is |
| $R$ | 2.5 $\mu\text{m}$ | Sphere radius |
| $r_{exo}$ | 50 nm | Exosome radius see Tab. 5 |
| $\Omega$ | 0.01, 0.03, 0.05 | Fraction of pole area particle starts/are recruited into, and exocytosis occur in |
| $\alpha$ | 0.2, 0.5, 0.8 | Mobile membrane fraction, see Tab. 5 |
| $k_{rec}$ | 60/s | Particle recruitment rate |
| $k_{off}$ | 1.0, 0.5, 0.05 /s | Particle dissociation rate |
| $D_m$ | 0.045, 0.0045, 0.00045 $\mu m^2/s$ | Particle diffusion rate |

#### Simulation details

To simulate the Cdc42 and septin ring models, we use a Finite Element Method (FEM) solver in space and finite differences solver in time using the FEniCSx software [Logg et al., 2012]. Following the logic in [Borgqvist et al., 2021], a mixed implicit-explicit Euler scheme is used in time. The non-linear reaction terms are treated as explicit, while all linear terms and gradients are treated as implicit. This approach incorporates non-linear terms into the load vector, allowing the FEM weights *ξ* at time *t* to be obtained by solving a large linear system. Since this linear system must be solved at each time step, the choice of linear solver is crucial for simulation performance. Given that our linear system is non-symmetric, we use the GMRES (Generalized Minimal Residual Method) solver with incomplete LU factorization (ILU) as a preconditioner.

All models comprise coupled partial differential equations (PDEs) and ordinary differential equations (ODEs). We adopt a mixed-stepping approach, where: i) the PDEs are updated using the schema above, and ii) the ODEs are then updated using a third-order Runge-Kutta method (explicit midpoint method). To ensure solution correctness and reduce runtime, we employ an adaptive time-step algorithm. Additionally, to guarantee accurate solutions, we use a finite-element mesh generated in the Gmsh software [Geuzaine and Remacle, 2009] with a high node density of 9451 nodes.

Cluster area in the model is computed as in [Borgqvist et al., 2021]; by computing the fraction of nodes that fulfill |*Cdc42_{max} – Cdc42*| *< 0.2 (Cdc42_{max} – Cdc42_{min})*. Septin ring diameter is computed using the algorithm in Fig. 5d, where the points constituting the septin ring are selected by filtering out all nodes where |*P_{max} – P*| *> 0.2 P_{max}* (results are robust to different threshold values than 0.2). Cluster area and ring diameter are computed at steady-state (after the model has been simulated for a long time), when there is only a single Cdc42-GTP cluster. This avoids measuring the area for multiple clusters and ensures that transient model dynamics are not measured for more fair comparison.

#### Computing septin ring diameter

The simulation produces a set of coordinates (points) that correspond to the septin ring, see Fig. 5d. For each inner point (dmin – blue Fig. 5d) the inner diameter is computed, and then the outer diameter is computed (dmax – green Fig. 5d) for each outer point. Then the mean of all dmax and dmin is computed (right part Fig. 5d).

### Statistics and reproducibility

All measurements are based on at least two independent biological replicates. Results from individual replicates were compared, and no major differences were noted.

## Data availability

Yeast strains as well as microscopy raw files are available upon reasonable request. Source data are provided together with this paper.

## Code availability

Additional information on image analysis approaches described in “Methods” and previous publications is available upon reasonable request. The code for model simulations can be found here: https://github.com/sebapersson/cdc42_and_septin_ring_paper.

## Acknowledgments

We thank Nils Johnsson and Daniel Lew for sharing yeast strains and feedback on the manuscript, and Michael Lanz and Jan Skotheim for sharing proteomics data. This work was supported by the Swedish Research Council (VR2017-05117 and VR2023-04319), the Swedish Foundation for Strategic Research (FFL15-0238) to MC, the Human Frontier Science Program (career development award to K.M.S.), the Deutsche Forschungsgemeinschaft (DFG, German Research Foundation) through SFB 1064 (Project-ID 213249687) and SFB 1309 (Project-ID 325871075) to R.S., and the Helmholtz Gesellschaft (K.M.S. and R.S.). The funders had no role in study design, data collection and analysis, decision to publish, or preparation of the manuscript.

## Declaration of interests

The authors declare no competing interests.

## Notes

### Competing Interest Statement

The authors have declared no competing interest.

