## Supplemental Information for "The origin of septin ring size control in budding yeast"

**Table S1. List of strains used in the study.**

| # | Strain name | Genotype | Strain background | Origin | Figure |
|---|---|---|---|---|---|
| 1 | DLY16570 | *Mat a/α; rsr1::TRP1/rsr1::TRP1 BEM1-GFP:LEU2/BEM1-GFP:LEU2 GIC2(1-208)-tdTomato:KAN$^{R}$/GIC2* | YEF473 | Kuo et al., Current Biology, 2014 | 1e,g, 2e, S6b |
| 2 | DLY16571 | *Mat a/α; CDC24$^{38A}$/CDC24$^{38A}$ rsr1::TRP1/rsr1::TRP1 BEM1-GFP:LEU2/BEM1-GFP:LEU2 GIC2(1-208aa)-tdTomato:KAN$^{R}$/GIC2* | YEF473 | Kuo et a.l, Current Biology, 2014 | 1e,g, 2e, S6b |
| 3 | KSY195-1 | *Mat α; ADE2 cdc10::CDC10-mCitrine-ADH1term-HIS3* | W303 | Kukhtevich et al., Nature Communications, 2020 | 3a-e |
| 4 | KSY260-2 | *Mat α; ADE2 cdc10::CDC10-mCitrine-ADH1term-HIS3 bni1Δ::KlacURA3* | W303 | Kukhtevich et al., Nature Communications, 2020 | 3a-e |
| 5 | KSY274-1 | *Mat α; ADE2 cdc10::CDC10-mCitrine-ADH1term-HIS3 bnr1Δ::KlacURA3* | W303 | this study | 3a-e |
| 6 | KSY277-2 | *Mat α; ADE2 cdc10::CDC10-mCitrine-ADH1term-HIS3 spa2Δ::KlacURA3* | W303 | this study | 3a-e |
| 7 | KSY278-1/2 | *Mat α; ADE2 cdc10::CDC10-mCitrine-ADH1term-HIS3 bud6Δ::KlacURA3* | W303 | this study | 3a-e |

| | | | | | |
|---|---|---|---|---|---|
| 8 | KSY306-3 | *Mat a; his3::LexA-ER-AD-TF-HIS3 whi5::kanMX6-LexApr-WHI5-ADH1term-LEU2 exo84::Exo84-mCitrine-ADH1term-cglaTRP1, cdc10::CDC10-mNeptune2.5-ADH1term-ADE2* | W303 | this study | 4d-g |
| 9 | KSY309-7/8 | *Mat a; his3::LexA-ER-AD-TF-HIS3 whi5::kanMX6-LexApr-WHI5-ADH1term-LEU2 exo84::Exo84-mCitrine-Adh1term-cglaTRP1, cdc10::CDC10-mNeptune2.5-Adh1term-ADE2 bni1∆::NatMX6* | W303 | this study | 4d-g |
| 10 | KSY317-9/12 | *Mat a; his3::LexA-ER-AD-TF-HIS3 whi5::kanMX6-LexApr-WHI5-ADH1term-LEU2 abp140::Abp140-mCitrine-Adh1term-cglaTRP1 cdc10::CDC10-mNeptune2.5-Adh1term-ADE2* | W303 | this study | 4a-c |
| 11 | KSY318-2 | *Mat a; his3::LexA-ER-AD-TF-HIS3 whi5::kanMX6-LexApr-WHI5-ADH1term-LEU2 abp140::Abp140-mCitrine-ADH1term-cglaTRP1 cdc10::CDC10-mNeptune2.5-ADH1term-ADE2 bni1∆::NatMX6* | W303 | this study | 4a-c |
| 12 | KSY342-1/2 | *Mat a/α; rsr1::TRP1/rsr1::TRP1 bem1::BEM1-GFP-LEU2/bem1::BEM1-* | YEF473 | this study, based on Kuo et al., | S3 |

| | | | | | |
|---|---|---|---|---|---|
| | | *GFP-LEU2 GIC2/gic2::GIC2(1-208)-tdTomato:KAN$^{R}$ his3/his3::LexA-ER-AD-TF-HIS3 WHI5/whi5::LexApr-WHI5-ADH1term-URA3* | | Current Biology, 2018 | |
| 13 | KSY343-2 | *Mat a/α; CDC24$^{38A}$/CDC24$^{38A}$ rsr1::TRP1/rsr1::TRP1 bem1::BEM1-GFP-LEU2/bem1::BEM1-GFP-LEU2 GIC2/gic2::GIC2(1-208)-tdTomato:KAN$^{R}$ his3/his3::LexA-ER-AD-TF-HIS3 WHI5/whi5::LexApr-WHI5-ADH1term-URA3* | YEF473 | this study, based on Kuo et al., Current Biology, 2019 | S3 |
| 14 | STY892 | *Mat a; BEM1::BEM1-GFP TRP1, CDC11::CDC11-mCherry URA3, CDC24::CDC24K525E, K801A-natNT2* | JD47 | Chollet et al., Journal of Cell Science, 2020 | 5h-j |
| 15 | STY893 | *Mat a; BEM1::BEM1-GFP TRP1, CDC11::CDC11-mCherry URA3, CDC24::CDC24-natNT2* | JD47 | Chollet et al., Journal of Cell Science, 2020 | 5h-j |
| 16 | Y1818, Y1819 | *Mat a/α; rsr1::TRP1/rsr1::TRP1 BEM1-GFP::LEU2/BEM1-GFP::LEU2 CDC10/cdc10::CDC10-mCherry-ADH1term-KanMX* | YEF473 | this study, based on Kuo et al., Current Biology, 2014 | 6a,b, S6 |
| 17 | Y1820, Y1821 | *Mat a/α; CDC24$^{38A}$/CDC24$^{38A}$ rsr1::TRP1/rsr1::TRP1 BEM1-* | YEF473 | this study, based on Kuo et al., Current Biology, 2016 | 6a,b, S6 |

| | | |
|---|---|---|
| | | *GFP:LEU2/BEM1-GFP:LEU2 CDC10/cdc10::CDC10-mCherry-ADH1term-KanMX* |

**Septin ring formation model based on previously described principles is not robust to parameter changes**

Our results suggest that exocytosis is diffused in *bni1*Δ. Since exocytosis displaces proteins from the polarity site in *S. pombe* [Gerganova et al., 2021] and has been proposed to sculpt the septin ring in *S. cerevisiae* by displacing septin [Okada et al., 2013], we hypothesized that diffused exocytosis could produce a larger ring diameter by septin displacement (Fig. 4h). To evaluate the plausibility of this, we turned to computational modeling.

Okada et al. [Okada et al., 2013] showed via computational modeling that exocytosis can sculpt a septin ring. Based on this suggested mechanism, we implemented a model. However, even though this model was able to form a ring, it was not robust to changes in model parameters including cell volume. Additionally, more recent studies [Lai et al., 2018] have shown that septin ring formation can start in the absence of a Sec4 signal (exocytosis marker), suggesting additional mechanisms in addition to exocytosis-promoting ring formation.

Therefore, we implemented a new model (Fig. S4a, b). The precise recruitment mechanism of septin is unknown, but septin monomers interact with polarity proteins Gic1/2, Axl2, and Cdc24 [Iwase et al., 2006; Chollet et al., 2020; Kang et al., 2024], hence we model that Axl2 (representing the other polarity factors) recruits septin. Cdc24 has been observed to bind Cdc11 in the Cdc42-GTP cluster, but not in the septin ring [Chollet et al., 2020]. To achieve ring formation without exocytosis, we model that polarity proteins (here Axl2) bind to septin and inhibit polymerization within the Cdc42 cluster area, subsequently forcing polymerization at the cluster periphery (Fig. S4b). Lastly, following Okada et al. [Okada et al., 2013], exocytosis is directed towards the pole where it displaces proteins, and septin recruits GAP proteins (Fig. S4b).

Because septin recruitment is likely cell-cycle triggered [Lai et al., 2018], and to reduce simulation time (one simulation takes > 100 hours), we initiated simulations from a Cdc42-GTP cluster (Fig. S4c). As observed experimentally [Okada et al., 2013], also in the model Cdc42-GTP concentration initially decreased, and then increased when a stable ring formed. In the Okada et al. [Okada et al., 2013] model this was partially achieved by having septin act as a diffusion barrier. Interestingly, in our model without any diffusion barrier, the septin-recruited GAP achieved a similar effect by trapping and concentrating Cdc42-GTPs within the ring (Fig. S4c).

Next, we investigated if diffused exocytosis produced a larger ring diameter. Starting from the same Cdc42-GTP cluster we simulated 10 cells for two model parameter combinations. Exocytosis was modeled to be directed towards the Cdc42 cluster, and as we use a finite element method (FEM) simulator where the model's geometry is meshed, we simulated concentrated and diffused exocytosis by allowing vesicles to hit approximately 0.03% or 0.34% of the mesh nodes respectively. Using a custom ring diameter computing algorithm (Fig. 5d), we found that the septin ring diameter was unaffected by diffused exocytosis (Fig. S4d).

The lack of effect of diffused exocytosis on ring diameter can depend on model parameter values. Parameters are tuned to yield realistic dynamics, however, due to simulation runtime (a single run takes >100 hours) an exhaustive search is infeasible. Thus, we created a simple fast-to-simulate particle model that captures key properties of ring formation. This includes that septin particles are recruited into a cluster area (representing the Cdc42-GTP cluster), and that the septin particles diffuse on the membrane, recycle, and are displaced by exocytosis (Fig. S5). Surprisingly, for different model parameters, diffused exocytosis did not produce a large septin ring diameter (Fig. S5). Similar results hold for a simpler particle simulator, where non-moving particles are displaced solely by exocytosis (Fig. S5).

In summary, the septin ring formation model suggested in Okada et al. [Okada et al., 2013] is not robust to changes in cell volume, while our model partially based on the mechanism suggested in Okada et al. does not explain why diffused exocytosis leads to an increase in septin ring diameter.

### Supplemental figures

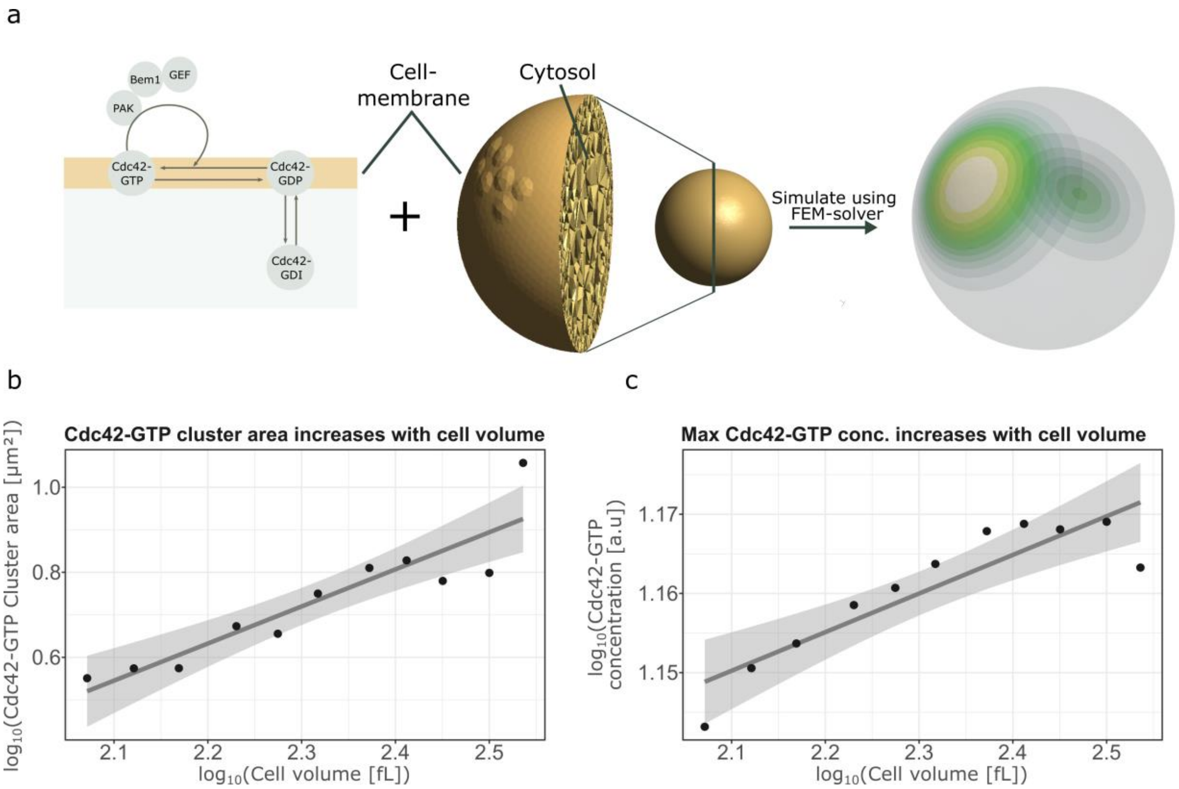

**Figure S1. Alternative computational model of Cdc42 polarization with positive feedback.** (a) Schematic drawing of the alternative model with positive feedback from [Borgqvist et al., 2021] (left), and result from a representative simulation (right). (b) Cdc42-GTP cluster area measured at steady-state (after long simulation time) for the model plotted against cell volume in double logarithmic scale. In each case, n=11 cells with volumes ranging from 115 to 345 fL were simulated starting from random initial conditions. Note that the volume interval differs from Fig. 1 as the model polarizes in a different parameter regime compared to the models in Fig. 1. (c) Cdc42-GTP maximum concentration against cell volume for the same simulated cells as in panel b, measured at the same time-point as the cluster area. Solid lines in (b-c) show linear regression fits.

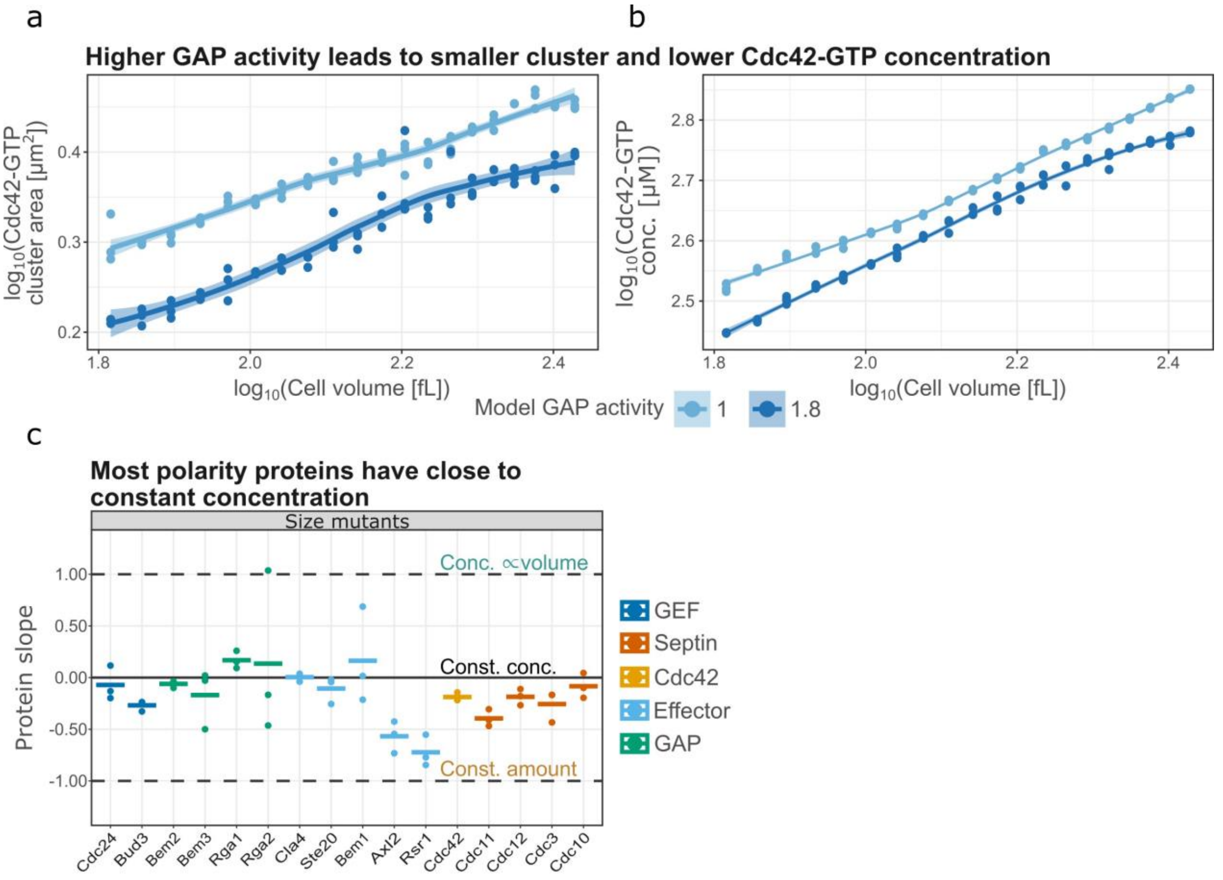

**Figure S2. Increased GAP activity and protein dilution in the positive feedback model.** (a-b) Cdc42-GTP cluster area (a) and Cdc42-GTP concentrations (b) for the positive feedback model measured at steady-state (after long simulation time) plotted against cell volume in double logarithmic scale for normal (1) and stronger (1.8) GAP activity. The increased GAP activity mimics negative feedback by reducing Cdc42 activation. In each case, n=60 (three replicates per volume) cells with volumes ranging from 65 to 270 fL were simulated starting from random initial conditions. Solid lines show loess smoothings. (c) Protein slope as in Fig. 2, but obtained from cell size mutants (see [Lanz et al., 2024] for more details). Bars show the mean value of n=3 biological replicates.

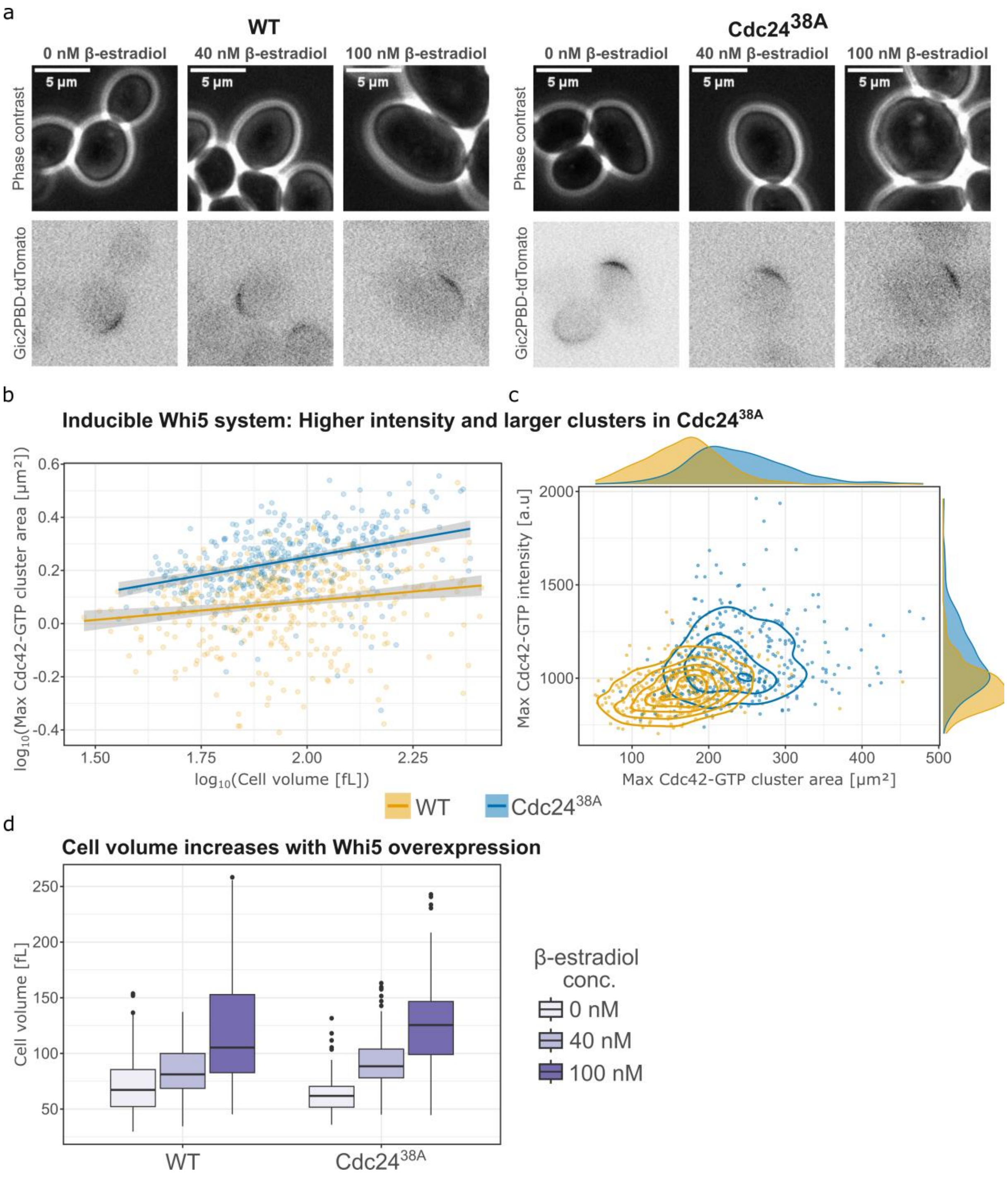

**Figure S3. Cdc42-GTP cluster scaling and intensity in strains with hormone inducible Whi5.** (a) Representative microscopy images of budding yeast cells (phase contrast) and Cdc42-GTP clusters (Gic2PBD-tdTomato) for wild-type Cdc24 (WT) and Cdc24$^{38A}$ cells. (b) Maximum Cdc42-GTP cluster area plotted against corresponding cell volume in double logarithmic scale. Solid lines show linear regression fits. (c) Maximum Cdc42-GTP intensity in the cluster at the time where the cluster reaches its maximum area, plotted against the corresponding Cdc42-GTP cluster area. (d) Cell volume for wild-type Cdc24 and Cdc24$^{38A}$ cells for different β-estradiol concentrations inducing Whi5 expression. In each plot, wild-type: 0 nM β-estradiol n = 123, 40 nM β-estradiol n = 111, 100 nM β-estradiol n = 181; Cdc24$^{38A}$: 0 nM β-estradiol n = 122, 40 nM β-estradiol n = 99, 100 nM β-estradiol n = 145.

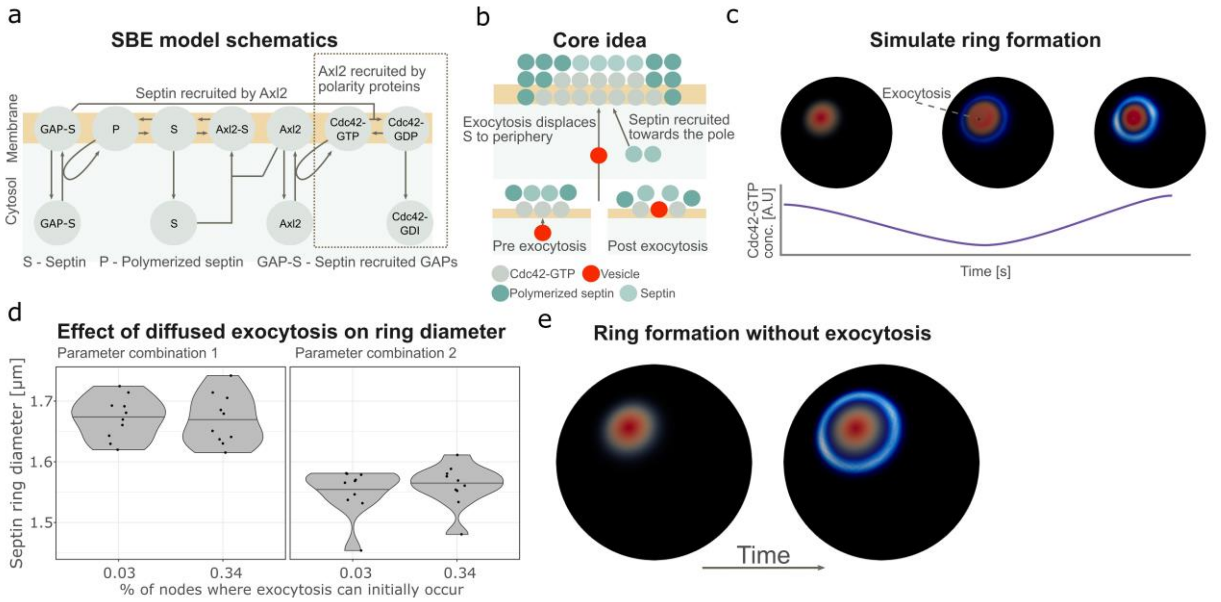

**Figure S4. The *Septin Binding and Exocytosis* (SBE) model does not explain why septin ring size increases with diffused exocytosis.** Model reaction schematics (a) and core idea (b) of the SBE model. Briefly, septin is recruited by Axl2, and on the membrane septin binds to Axl2. This binding prevents polymerization in the cluster center, which promotes polymerization and, subsequently, ring formation at the cluster periphery. Additionally, exocytosis is directed towards the cluster, which further pushes septin to the periphery. (c) Representative example showing Cdc42 polarization (red) and consecutive septin ring formation (blue). Cdc42-GTP concentration first decreases when septin is recruited, and then increases when a stable septin ring starts to form. (d) Septin ring diameter (*dmin+dmax)/2* (see Fig. 5) for the SBE model plotted for two parameter combinations and two levels of diffused exocytosis. For each condition, n=10 simulations, all starting from the same Cdc42-GTP cluster, were performed. In each case, the model was simulated for a long time to reach a stable ring, and then septin ring diameter was measured. The number of nodes that can be hit corresponds to nodes where the concentration of Cdc42 fulfills: *Cdc42-GTP > ε*max(Cdc42-GTP)*, where a smaller *ε* corresponds to more diffused exocytosis. (e) Example of septin ring formation for the SBER model without exocytosis. Note that without exocytosis, the SBER model is equivalent to the SBE model.

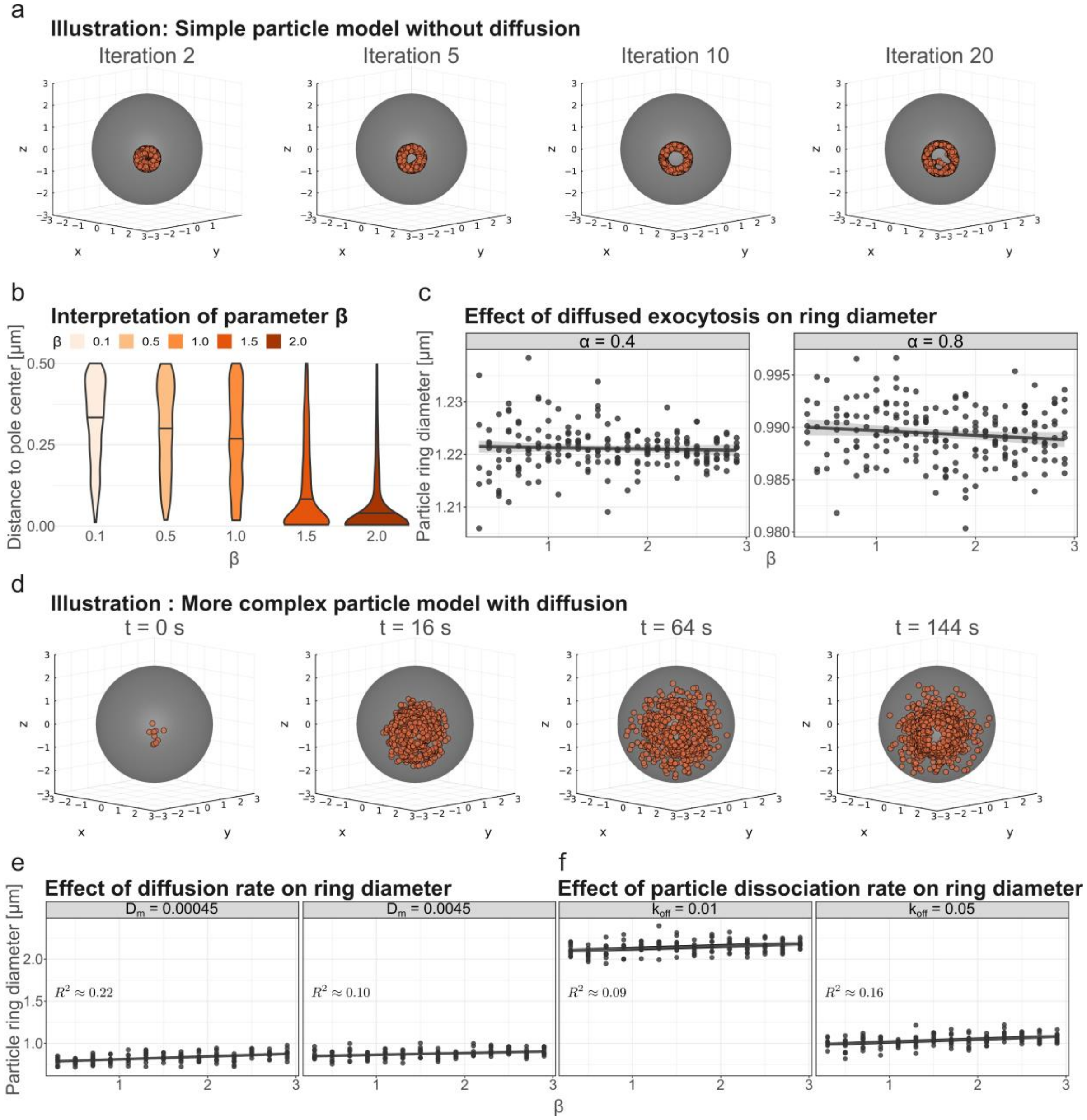

**Figure S5. Particle models suggest that diffused exocytosis itself does not produce a larger septin ring diameter.** (a) Illustration of the simple particle model without diffusion. Simulations start with particles covering 3% of a circular cluster area (red), followed by 20 iterations of exocytosis. (b) Parameter β affects the spread of exocytosis. Exocytosis can occur in the same circular cluster area covering 3% of the surface that particles start in or are recruited into. For a large β, exocytosis is concentrated, as seen by the distribution of the distance to the cluster area from where exocytosis occurs. Meanwhile, for a smaller β, exocytosis is more diffused. (c) Septin ring diameter as a function of β for different values of the exocytosis model parameter α (see methods). Each plot shows n=210 simulations. (d) Illustration of a more complex particle model than in contrast to the earlier particle model also includes diffusion and particle recruitment, where particles are recruited into a cluster occupying 3% of the area, and then diffuse at rate $D_m$, disassociate at rate $k_{off}$, and undergo exocytosis. (e-f) Septin ring diameter as a function of β for the second particle model, shown for different values of the diffusion rate $D_m$ (e) and the dissociation rate $k_{off}$ (f). Low diffusion rates and high dissociation rates lead to larger rings with concentrated exocytosis. Each plot shows n=140 simulations. (c,e-f) Solid lines show linear regression fits.

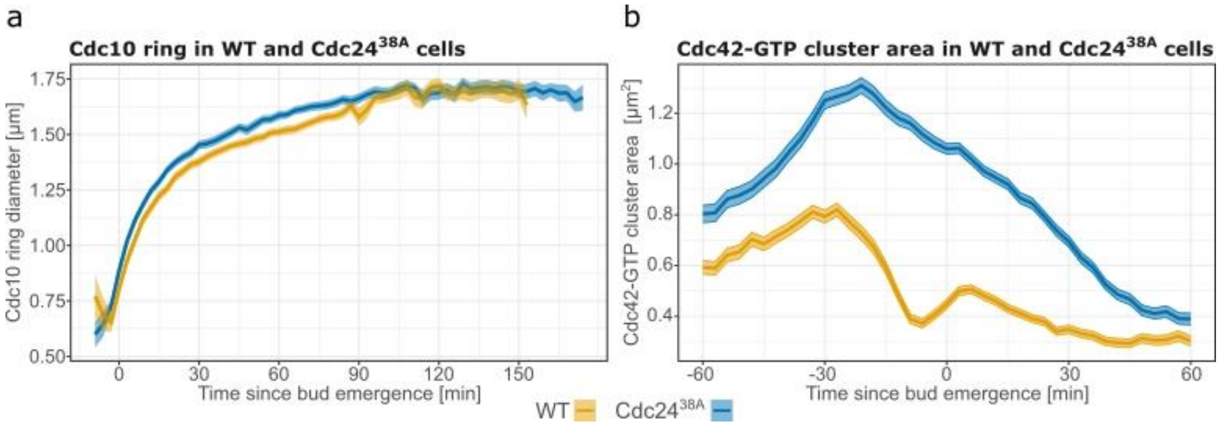

**Figure S6. Septin ring and Cdc42-GTP cluster formation in negative feedback mutant.** (a) Mean Cdc10 ring diameter measurements from microscopy images plotted over the cell cycle for WT (n = 68 cells) and Cdc24$^{38A}$ (n = 101 cells). (b) Mean Cdc42-GTP cluster area measurements from microscopy images plotted over the cell cycle for WT (n = 453 cells) and Cdc24$^{38A}$ (n = 465 cells). Single cell traces are aligned at bud emergence. The ribbons correspond to the standard error.
